# Efficient globin production during terminal erythropoiesis depends on the synergistic action of TENT5C poly(A) polymerase and LARP4/5

**DOI:** 10.1101/2024.11.14.623596

**Authors:** Michał Mazur, Natalia Gumińska, Aleksandra Brouze, Dominik Cysewski, Katarzyna Mleczko-Sanecka, Marta Niklewicz, Monika Kusio-Kobiałka, Andrzej Dziembowski

## Abstract

Red blood cell development is a unique process where reduced transcriptome and proteome complexity facilitates vast hemoglobin production. Here, we describe the cooperative role of cytoplasmic poly(A) polymerase TENT5C and the poly(A) tail-protecting LARP4/5 RNA-binding proteins in ensuring proper hemoglobin production. TENT5C catalytic mutant knock-in mice display microcytic hypochromic anemia resembling constitutive knockout. TENT5C counteracts gradual globin mRNA deadenylation during erythropoiesis. In the late stages, TENT5C dysfunction leads to globin poly(A) tail shortening and a drastic reduction of mRNA levels in reticulocytes. Proteomic experiments revealed transient but specific association of TENT5C with LARP4/5. Indeed, LARP4/5 depletion leads to downregulation and poly(A) tail shortening of globin mRNAs. Furthermore, lack of TENT5C catalytic activity is accompanied by compensatory upregulation of LARP4/5. Finally, the importance of precise regulation of globin poly(A) tails by deadenylation and re-adenylation is highlighted by the destabilization of TENT5C by CCR4-NOT deadenylase complex-associated E3 ubiquitin ligase CNOT4.

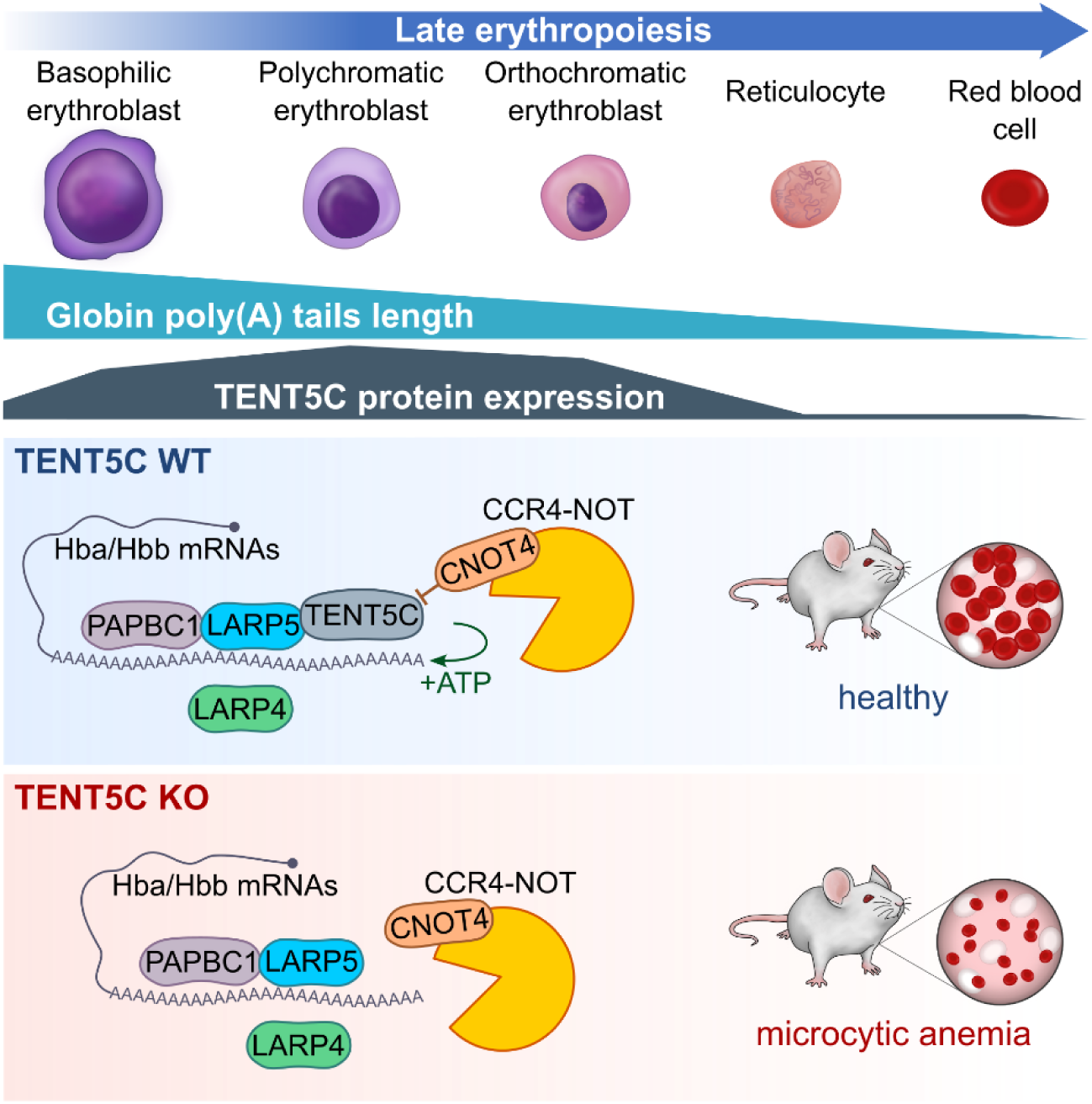

## INTRODUCTION

Every day, our bodies generate and break down approximately 200 billion red blood cells (RBCs), the only cells whose proteome is composed predominantly (∼98%) of hemoglobin^1–4^. Erythropoiesis spans the developmental pathway from stem cells to mature RBCs^5^. It begins with the differentiation of a hematopoietic stem cell (HSC) into the first committed erythroid progenitors burst-forming unit-erythroid (BFU-E) followed by colony-forming unit-erythroid (CFU-E). CFU-E progenitors divide 3 to 5 times and differentiate into proerythroblast (proE) followed by the basophilic erythroblast (basoE), polychromatophilic erythroblast (polyE), and orthochromatic erythroblast (orthoE), which exits the cell cycle. After the orthochromatic erythroblast extrudes the nucleus, it becomes a reticulocyte (retc), which finally matures into RBC^5–8^. Classical erythropoiesis occurs in the bone marrow (BM) and is organized around macrophage-centered aggregates called erythroblastic islands (EBIs). Here, erythroblasts are maintained by a central macrophage and are gradually maturing until enucleation stage, ultimately being released to the bloodstream^8–11^. During mouse embryonic development, the main reservoir for erythropoiesis is the fetal liver, where erythroid cells make up ∼90% of cellular population between E12.5 and E15.5^12–16^. In response to anemia or blood loss, the organism may compensate for RBC deficiency by triggering stress erythropoiesis, often termed extramedullary erythropoiesis, primarily occurring in the spleen or the liver^17–19^. This adaptive mechanism adjusts the RBC production according to physiological demand, regulated by various cytokines acting on EBIs. During adult erythropoiesis, the pro-inflammatory cytokines play an important regulatory role as regulatory factors maintaining homeostasis^9,20^. Regulatory cytokines, including TGF-β1, IL6, IFN-β, TPO, TSLP, or EPO, directly affect RBCs differentiation, and elevated levels of these cytokines can induce stress erythropoiesis while impairing steady-state erythopoiesis^17,21–26^.

Intracellular changes during RBC development are defined by the degradation of most cellular components alongside the continuous production and accumulation of hemoglobin^3,4,6^. In terminal erythropoiesis, active RNA polymerase II is mostly allocated to erythroid-specific genes^27^. Accordingly, maturing erythroblast transcriptomes consist mainly of globin mRNAs, and their complexity is further reduced as cells differentiate^28^. Autophagic degradation is indispensable for the clearance of organelles, while proteome reduction is maintained by degradation in the proteasome^29,30^. However, the most drastic change for cells during terminal erythropoiesis is enucleation, which physically restricts the production of nascent transcripts, leaving residual mRNAs to be translated or degraded^31–34^.

Because of its uniqueness, the regulation of mRNA metabolism during RBC development has been extensively studied. Previous reports focused on translational control, splicing regulation, or mRNA degradation^35,36^. However, despite the importance of poly(A) tails in protein synthesis and the influence of deadenylation rates on mRNA stability, little is known about the regulation of poly(A) tails of globin mRNAs. Moreover, the mechanisms sustaining globin mRNA translation and stability after enucleation remained unclear. In this respect, some reports were suggesting protection from deadenylation by RNA-binding protein αCPs (PCBP1 and PCBP2) specifically for α-globin mRNA^37,38^.

Herein, we studied the role of cytoplasmic poly(A) polymerase TENT5C in RBC development. We demonstrate that TENT5C re-adenylates globin mRNAs, which is essential for the terminal stages of RBC maturation. In murine erythroblasts expressing TENT5C-catalytically inactive mutant, protein production of globins is deregulated and their corresponding mRNAs drop significantly in reticulocytes, leading to microcytic anemia compensated by stress erythropoiesis in the spleen. We propose a mechanism in which RNA-binding proteins LARP4/5 cooperate with TENT5C to protect globin mRNA poly(A) tails and facilitate TENT5C-driven polyadenylation. Furthermore, we show that TENT5C protein levels are tightly and dynamically controlled. Specifically, we identify TENT5C as highly unstable due to its rapid degradation by the CCR4-NOT deadenylase complex-associated E3 ubiquitin ligase CNOT4.

## RESULTS

### TENT5C catalytic activity is required for normal erythropoiesis

To gain insight into the mechanism of TENT5C-mediated regulation of erythropoiesis, we first aimed to characterize the phenotype associated with TENT5C dysfunction in mice. To assess if the catalytic activity of TENT5C plays a direct role in erythropoiesis, we analyzed the catalytically dead TENT5C (D90N; D92N) knock-in mice (*Tent5c^cat^*), which turned out to be indistinguishable from a constitutive knock-out (*Tent5c^ko^)*^39^. Whole blood analysis revealed abnormal parameters with decreased mean corpuscular hemoglobin, mean corpuscular hemoglobin concentration (MCH, MCHC), and mean corpuscular volume (MCV) of RBCs, pointing to microcytic hypochromic anemia in *Tent5c^cat^* mice (Fig. 1A). Moreover, catalytic mutants showed elevated counts of RBCs and platelets (Fig. 1A) with no difference in white blood cells and plasma albumin (Fig. 1A), indicating the absence of immune response and renal/hepatic dysfunction.

**Figure 1.**
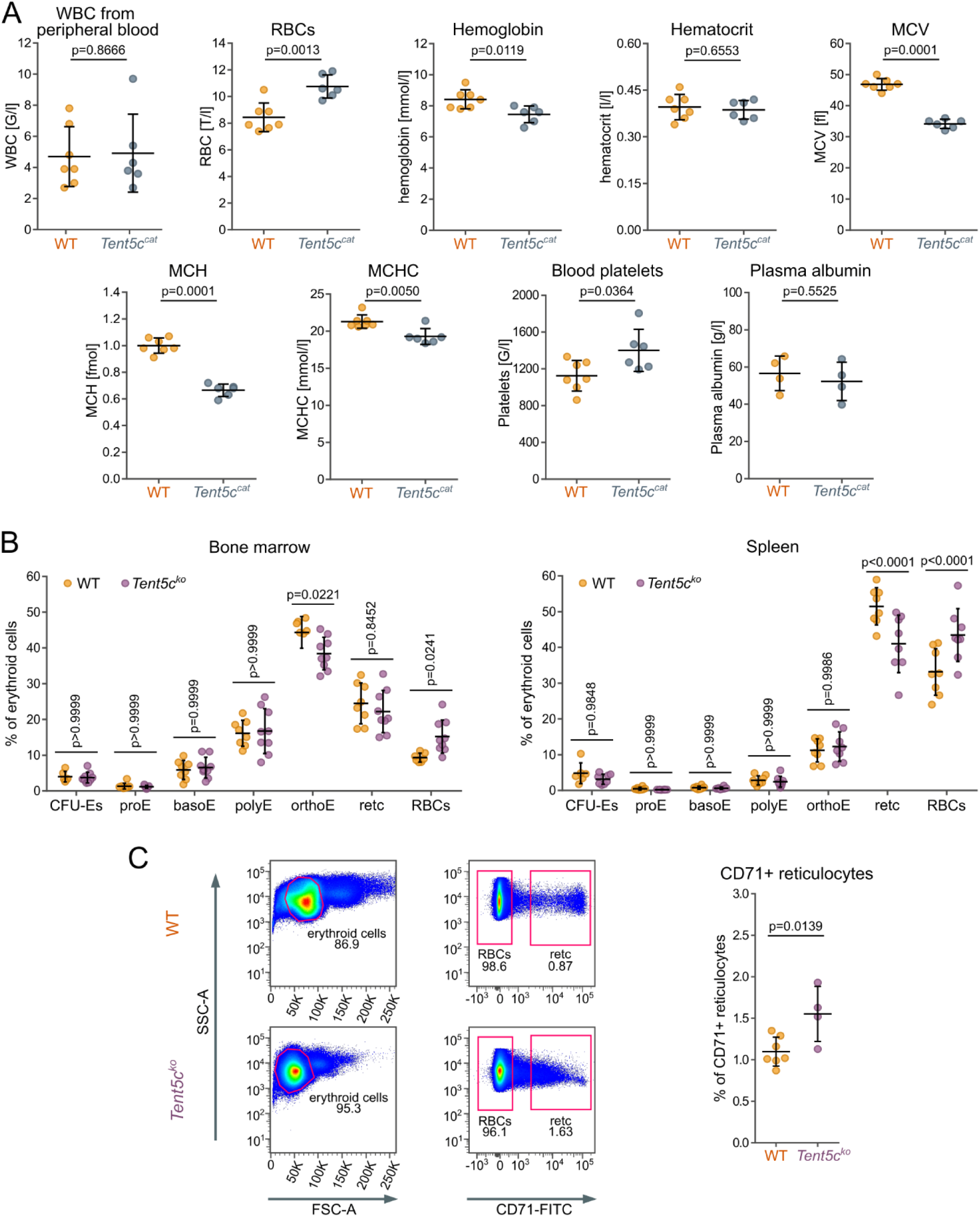
TENT5C catalytic activity is required for normal erythropoiesis. (A) Quantification of: white blood cells (WBC) isolated from peripheral blood, red blood cells count (RBCs), Hemoglobin, Hematocrit, mean corpuscular volume (MCV), mean corpuscular hemoglobin (MCH), mean corpuscular hemoglobin concentration (MCHC), blood platelets (WT *n*=7, *Tent5c^cat^ n*=6) and plasma albumins (WT *n*=4, *Tent5c^cat^ n*=4). Data presented as mean ± SD, and significance was assessed by t-Student test. (B) Quantification of erythroblast subpopulations in the bone marrow (left panel) and the spleen (right panel) (WT *n*=8, *Tent5c^ko^ n*=8). Data presents as mean ± SD, and significance assessed by two-way ANOVA. (C) Quantification of CD71-FITC positive reticulocytes in the blood (WT *n*=7, *Tent5c^ko^ n*=4). Data presented as mean ± SD, and significance was assessed by *t*-Student test.

Then, to evaluate the effect of *Tent5c^ko^* on RBC development, we performed a flow cytometry analysis of blood and erythropoietic organs responsible for definitive erythropoiesis (bone marrow, spleen, fetal liver). Animals with *Tent5c* deficiency displayed slightly altered levels of erythroid subpopulations, with no difference in less differentiated cells (CFU-Es, proE, basoE, and polyE). However, we observed significantly decreased levels of orthoE in the BM and reticulocytes in the spleen, which was accompanied by significantly increased levels of mature RBCs in both organs of *Tent5c^ko^* animals (Fig. 1B). Additionally, we measured the level of highly immature reticulocytes (CD71^+^) circulating in the bloodstream and found that *Tent5c^ko^* mice have ∼1.5-fold elevated levels of these cells (Fig. 1C). These observations suggest faster but compromised differentiation during definitive erythropoiesis of *Tent5c^ko^* animals, marked by elevated levels of highly immature circulating reticulocytes. In the fetal liver, at the E15.5 stage of embryonic development, alterations in *Tent5c^ko^* are analogous to adult erythropoiesis but apply to basoE and polyE (Fig. S1A, right panel), and they were not observed at the earlier stage of E14.5 (Fig. S1 A, left panel).

Next, to check whether TENT5C-dependent anemia exhibits symptoms of chronic inflammation, potentially regulating microenvironmental signals, we measured levels of different cytokines in the blood serum of *Tent5c^ko^* using cytometry bead arrays. Notably, several proinflammatory cytokines, including TPO, CXCL1, IL12p40, IL6, CCL5, TSLP, IFN-β, TGF-β1 and GM-CSF were upregulated in *Tent5c^ko^* in comparison to wild type (WT) (Fig. 2A). There were no significant differences in the level of TNF-α or SCF (Fig. 2B). Consistently, IL-10, IFN-α, and CXCL12, cytokines that can exhibit anti-inflammatory roles, were downregulated in *Tent5c^ko^* mice (Fig. 2C). Some of the elevated pro-inflammatory cytokines (TGF-β1, GMC-SF, TPO) together with erythropoietin (EPO) (Fig. 2A, D) also take part in the megakaryopoiesis indicating that in *Tent5c^cat^* mice, the process of thrombocytes production may also be affected, which is backed by elevated levels of platelets (Fig. 1A). Most importantly, the elevated levels of EPO and altered cytokine profiles in *Tent5c^ko^* point to stress erythropoiesis. Therefore, we analyzed potential differences within EBIs in the bone marrow and spleens of adult mice by analyzing cellular aggregates. Indeed, *Tent5c^ko^* mice displayed significantly more aggregates corresponding to EBIs in the spleen compared to WT mice (Fig. 2E) with an opposite trend in the bone marrow (Fig. 2E). To verify this result, we checked the erythropoietic potential (EP) by measuring the number of CD71^+^Ter119^+^ erythroid progenitors in the bone marrow and spleens^17,20^ and observed downregulation of the EP in the bone marrow with a slight increase in the spleen (Fig. 2F). These findings are consistent with a well-established link between the pro-inflammatory signature of the serum cytokines and a shift of erythropoietic activity from the bone marrow to extramedullary sites, indicating stress hematopoiesis. Then, to elucidate in more detail the mechanisms responsible for changes in EBIs, we sought a correlation between studied cytokines and the expression levels of selected EBI macrophage receptors determined using flow cytometry. It has been shown that GM-CSF can disrupt EBIs in the bone marrow and downregulate CXCL12, CD163 and VCAM1 in their macrophages^40^. Indeed, we observed that *Tent5c^ko^* mice have increased levels of GM-CSF and decreased levels of CXCL12 in serum (Fig. 2A, 2C). Hence, this change explains lower expression levels of both CD163 and VCAM1 of EBI macrophages in the bone marrow based on their geometric Mean Fluorescence Intensity (gMFI). Interestingly, we did not observe the same changes in macrophages from the spleen EBIs (Fig. 2G, 2H). These changes, in concert with the pattern of cytokine levels in the serum, explain the observed shift towards stress hematopoiesis and the downregulation of EBIs in the bone marrow and their upregulation in the spleen.

**Figure 2.**
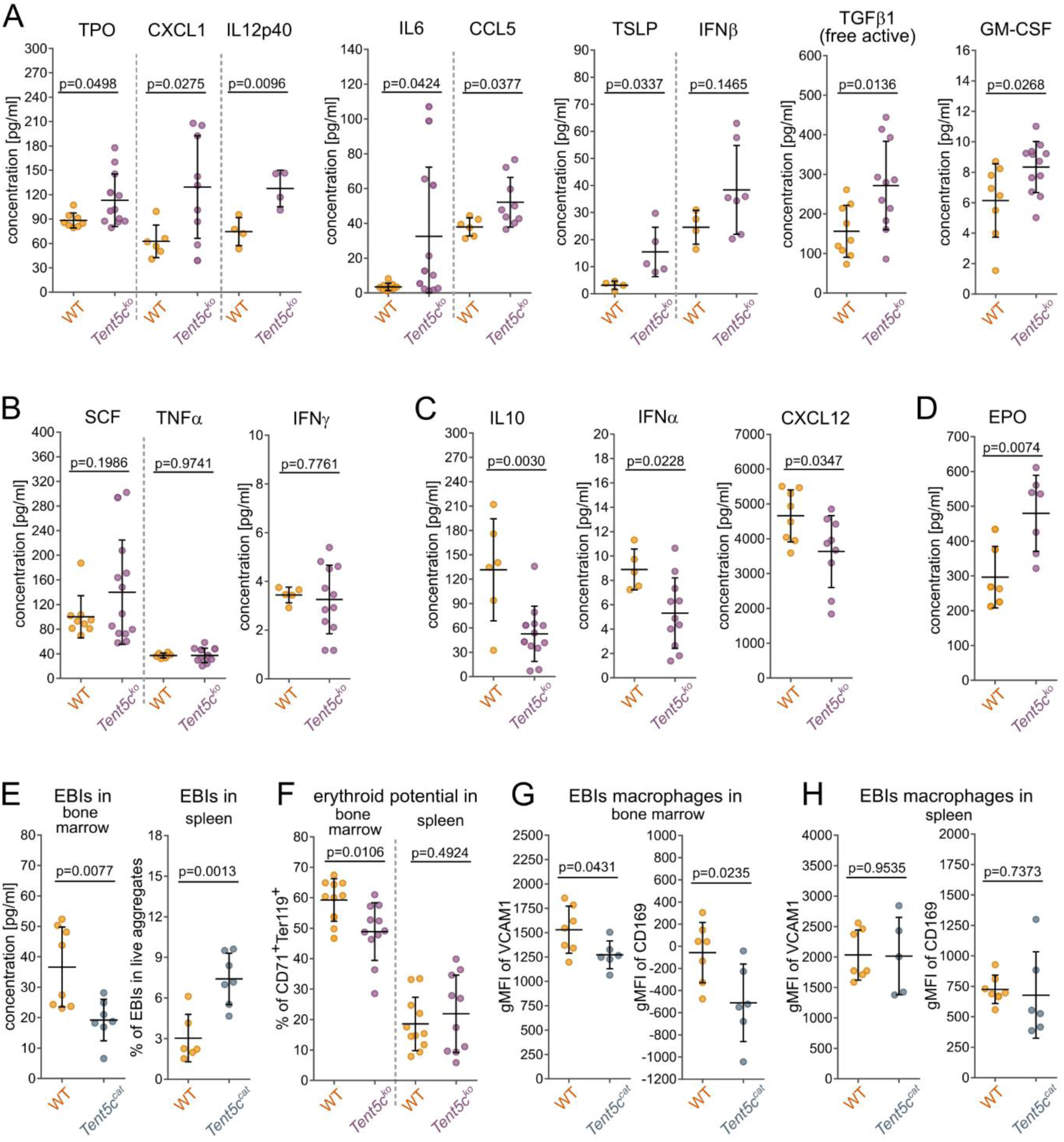
TENT5C inactivity induces splenic stress erythropoiesis. (A) Cytokines upregulated in *Tent5c^ko^* (WT *n*=6-9, *Tent5c^ko^ n*=6-12). Data presented as mean ± SD, and significance was assessed by *t*-Student test. (B) Cytokines with steady levels in *Tent5c^ko^* (WT *n*=5-9, *Tent5c^ko^ n*=12). Data presented as mean ± SD, and significance was assessed by *t*-Student test. (C) Cytokines downregulated in *Tent5c^ko^* (WT *n*=6-8, *Tent5c^ko^ n*=9-12). Data presented as mean ± SD, and significance was assessed by *t*-Student test. (D) Quantification of erythropoietin levels (WT *n*=6, *Tent5c^ko^ n*=7). Data presented as mean ± SD, and significance was assessed by *t*-Student test. (E) Percentages of EBIs in the bone marrow (left panel, WT *n*=8, *Tent5c^cat^ n*=7) and the spleen (right panel, WT *n*=6, *Tent5c^cat^ n*=7). Data presented as mean ± SD, and significance was assessed by *t*-Student test. (F) Quantification of erythroid potential showed as percentages of CD71^+^Ter119^+^ cells in the bone marrow and the spleen (WT *n*=10, *Tent5c^ko^ n*=11). Data presented as mean ± SD, and significance was assessed by *t*-Student test. (G) Quantification of VCAM1 and CD169 expression (gMFI) in EBI macrophages in the bone marrow (WT *n*=8, *Tent5c^cat^ n*=7). Data presented as mean ± SD, and significance was assessed by *t*-Student test. (H) Quantification of VCAM1 and CD169 expression (gMFI) in EBI macrophages in the spleen (WT *n*=6, *Tent5c^cat^ n*=7). Data presented as mean ± SD, and significance was assessed by *t*-Student test.

Finally, to assess the possibility of iron deficiency contributing to microcytic anemia of TENT5C-deficient mice, we have checked the iron levels in serum, spleen, and the liver, finding little to no effect in *Tent5c^cat^* mutants (Fig. S1B). In addition, we applied flow cytometry to assess the expression levels of iron uptake receptor CD71 from various erythroid tissues and found no significant effect in *Tent5c^ko^* mutants (Fig. S1C, S1D). All these results suggest that the anemic phenotype present in *Tent5c^ko^* and *Tent5c^cat^* mice is not iron-dependent.

We conclude that the lack of TENT5C catalytic activity is the main trigger of anemia in *Tent5c^ko^*/*Tent5c^cat^* mice leading to stress erythropoiesis. However, since the anemic phenotype originates from inactivity of an intracellular enzyme, it cannot be fully ameliorated by induced stress erythropoiesis, persisting through life of TENT5C-deficient mice.

### Globin mRNAs are primary TENT5C substrates in erythroblasts

To check if TENT5C can indeed be directly involved in hemoglobin synthesis, we characterized its expression pattern in different developmental stages. We took advantage of the *Tent5c-EGFP* knock-in mouse line, facilitating the estimation of TENT5C protein levels during cytometric analyses of erythroblasts from fetal livers and adult bone marrow.

The analysis of fetal livers from different developmental days (E12.5-E15.5) revealed the highest expression at E14.5 (Fig. 3A, left panel) with a peak in polyE (Fig. 3A, right panel). Similarly, for bone marrow, TENT5C-EGFP is highly expressed in late erythroblasts with a peak of expression in polyE, and downregulation in reticulocytes and RBCs (Fig. 3B). To corroborate our cytometric results with an orthogonal methodology, we performed an *ex vivo* culture of erythroid progenitors from the fetal livers and performed a western blot in a time course spanning 4 days of differentiation. Indeed, the highest TENT5C-EGFP expression was observed on day 2 of differentiation (Fig. S2A), where most cells correspond to the polyE differentiation stage.

**Figure 3.**
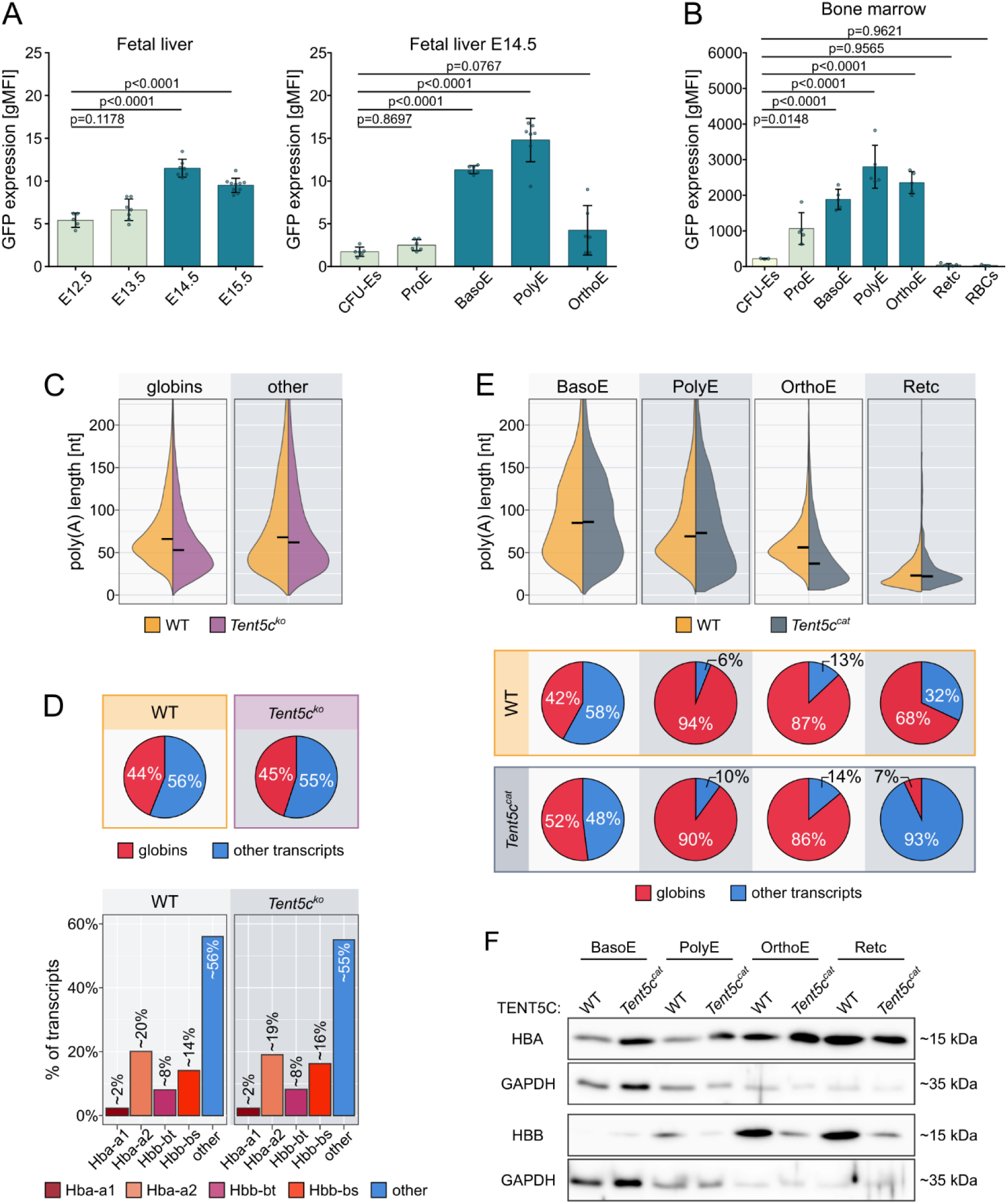
TENT5C counteracts globin mRNAs deadenylation during late erythropoiesis. (A) Quantification of TENT5C-EGFP expression (gMFI) in fetal liver erythroblasts at different developmental days E12.5-E15.5 (left panel) and quantification of TENT5C-EGFP expression (gMFI) in different subpopulations of E14.5 fetal liver erythroblasts (right panel) (*Tent5c-Egfp n*=5-10). Data presents as mean ± SD, and significance assessed by two-way ANOVA. (B) Quantification of TENT5C-EGFP expression (gMFI) in different subpopulations of erythroblasts from the bone marrow (*Tent5c-Egfp n*=5). Data presents as mean ± SD, and significance assessed by two-way ANOVA. (C) Distribution of poly(A) tail lengths in globin (left panel) and other (right panel) transcripts in DRS of E14.5 fetal livers (WT *n*=2, *Tent5c^ko^ n*=2). Black horizontal line indicates median. (D) Percentage of hemoglobins and other transcripts in DRS of E14.5 fetal livers. Upper panel (pie charts) aggregates all hemoglobins versus the other transcripts. Bottom panel (bar plot) show each hemoglobin separately versus the other transcripts (WT *n*=2, *Tent5c^ko^ n*=2). (E) Distribution of poly(A) tail lengths in globins during subsequent stages of erythropoiesis in cDNA seq (upper panel). Black horizontal line indicates median. The bottom panel (pie charts) show percentage of hemoglobins versus the other transcripts in examined cell subpopulations, respectively (WT *n*=3, *Tent5c^ca^*^t^ *n*=3). (F) Western Blot showing α– and β-globin protein abundances in subpopulations of erythroblasts from the BM acquired by FACS. Equal cell number loaded onto wells (5000 cells).

Since TENT5C is an enzyme directly catalyzing the addition of adenosines to poly(A) tails in the cytoplasm, we sought its mRNA substrates in erythroblasts using nanopore Direct RNA Sequencing (DRS), enabling transcriptome-wide poly(A) tail profiling^41^. To ensure enough starting material, we first used mRNA from the whole fetal livers of E14.5 developmental stage, during which we observed the highest TENT5C expression (Fig. 3A, left panel). Indeed, the globin transcripts were very abundant in the transcriptome, representing ∼45 % of reads in both WT and *Tent5c^ko^* mice (Fig. 3D, top panel), with a similar distribution of individual globins (Fig. 3D, bottom panel). Most importantly, globin mRNAs were the major transcript cluster with statistically significant shortening of poly(A) tails in the *Tent5c^ko^* fetal livers (Fig. 3C). The effect on globin poly(A) tails agrees with the microcytic anemia phenotype of *Tent5c^cat^* (Fig. 1A). Still, it contrasts with all the previously identified TENT5s substrates in other model systems, which were dominated by mRNAs possessing a signal peptide targeting them to the endoplasmic reticulum (ER)^39,42–45^. Surprisingly, we could not observe the difference in globin mRNAs abundances between WT and *Tent5c^ko^* fetal livers (Fig. 3D, top and bottom panel), suggesting a compensatory mechanism occurring in the heterogenous population of erythroblasts. Shortening of the poly(A) tails was not restricted to globin mRNAs, as we saw this effect in several other transcripts (Fig. 3C, S2B, Supplementary Table 1), some of which were recently described as putative targets of TENT5C^46^. However, we decided to focus on globin mRNAs as they were the most basic and abundant constituent necessary for hemoglobin production.

To precisely study the role of TENT5C in the regulation of globin mRNA poly(A) tail lengths dynamics during erythropoiesis, we sorted 4 cell populations (basoE, polyE, orthoE, retc) from the bone marrow of WT or *Tent5c^cat^* animals. We opted for Nanopore cDNA sequencing, which requires a much lower amount of starting material than DRS. The experiment was performed in triplicate. These datasets were even more dominated by globin reads than DRS from fetal liver. Therefore, we focused on globin mRNA exclusively. Generally, we could see a gradual shortening of the poly(A) tails during differentiation, with the shortest tails in reticulocytes represented by a median of only ∼19 A (Fig. 3E, top panel). When comparing WT to *Tent5c^cat^* erythroblasts throughout differentiation, we observed pronounced poly(A) tail shortening for all globin transcripts exclusively in orthochromatic erythroblasts (Fig. 3E, top panel), which is in line with our observation of the expression levels of TENT5C-EGFP (Fig. 3B). Notably, in the case of reticulocytes with very short tails, there is a dramatic drop in globin mRNA and protein abundances in *Tent5c^cat^* (Fig. 3E, bottom panel; Fig. 3F) without a visible effect on poly(A) tail length (Fig. 3E, top panel). This observation suggests that the primary role of TENT5C is stabilization of globin mRNAs, counteracting the decay mediated by a classic deadenylation-dependent pathway where decapping acts on transcripts with tails shorter than 20 A, which cannot bind poly(A) binding protein (PABP)^47,48^. Interestingly, at the earlier stages of differentiation, a distinct behavior dependent on the type of globin mRNA is observed. This points to transcriptional compensation, as nascent transcripts typically exhibit the longest poly(A) tails. It is particularly evident in the case of the *Hba-a1*, where the poly(A) tails are elongated in basoE and polyE *Tent5c^cat^* cells (Fig. S3A, S3B), corresponding with the upregulation of α-globin protein levels in these cells (Fig. 3F). In contrast, despite no visible effect on poly(A) tails or transcript abundance of *Hbb-bs or Hbb-bt* (Fig. S3A, S3B), we can observe a persistent downregulation of its protein levels already visible in polyE (Fig. 3F).

We concluded that TENT5C-mediated adenylation is critical to maintaining sufficient levels of globin mRNAs in reticulocytes. Additionally, TENT5C activity is crucial for proper β-globin protein production throughout terminal erythropoiesis.

### Proximity biotinylation reveals TENT5C interactome in erythroblasts

It has been demonstrated in different model systems that the substrate specificity of TENT5C is at least partially regulated by the attachment to the ER membrane, mediated by FNDC3A/B proteins^49^. However, globin mRNAs are not translated at the ER. Therefore, another targeting mechanism is expected to operate in erythroblasts. To identify erythroid-specific TENT5C interactors, we employed a modified TurboID proximity biotinylation assay followed by MS/MS analysis of fetal liver erythroblasts differentiated *ex vivo*. Again, we took advantage of *Tent5c-EGFP* knock-in mouse line and employed lentiviral vector expressing nanobody anti-EGFP fused with TurboID biotinylase^50^. By introducing this construct to either WT or *Tent5c-EGFP* erythroblasts, we were able to quantitatively determine the specificity of enriched proteins found in proximity to TENT5C-EGFP compared to the background found in the WT (Fig. 4A). This approach does not require the introduction of exogenous biotinylase-tagged TENT5C, which expression would be out of place in the earliest stages of erythropoiesis. Importantly, it was designed for the knock-in model in which the expression timing for TENT5C is in accord with the developmental timeline of erythroblasts. Hence, we expected the enriched proximal proteins to be physiologically relevant interactors of TENT5C.

**Figure 4.**
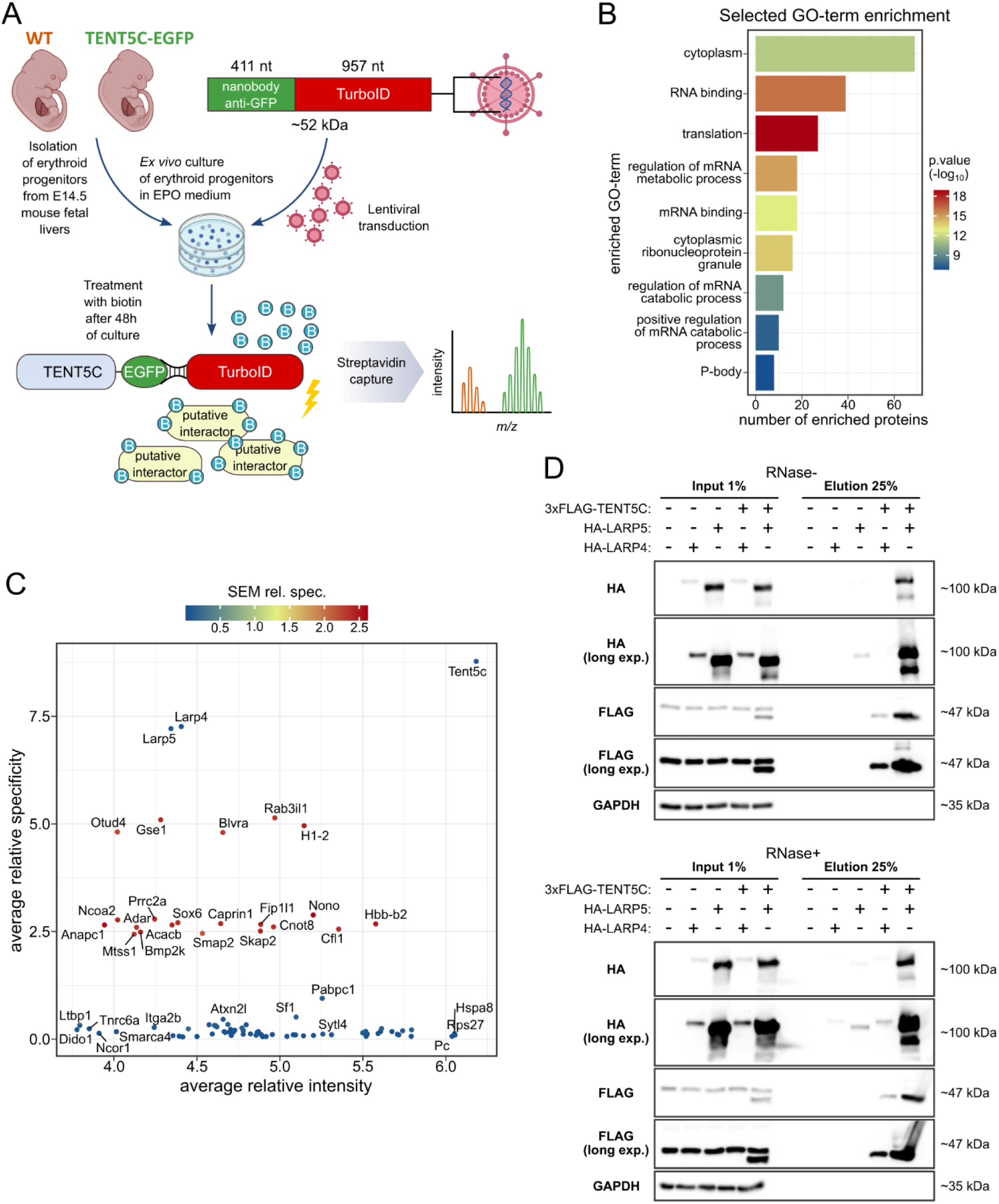
TENT5C protein interactome during erythropoiesis. (A) Schematic representation of the TurboID experiment. (B) Selected Gene Ontology (GO) terms for enriched group of proteins found in 3 biological replicates. Visualised by number of proteins falling into a category on X-axis, type of GO term on Y-axis, colored for –log10(p.value). (C) Scatterplot representing average relative specificity (Y-axis) and average relative intensity (X-axis) for enriched proteins found in overlap of 3 biological replicates. Datapoints colored for standard error of the mean (SEM). (D) Western Blot of CO-IP experiment in HEK293 cells after ectopic overexpression of HA-LARP4/5 and 3xFLAG-TENT5C. 1% inputs applied on gel (left side); 25% of elutions applied on gel (right side). In either fraction the lane order is the same and represents (from left to right): untransfected control, HA-LARP4 overexpression, HA-LARP5 overexpression, 3xFLAG-TENT5C and HA-LARP4 co-overexpression, 3xFLAG-TENT5C and HA-LARP5 co-overexpression. Top panel-lysates were not subjected to RNAse treatment prior to CO-IP. Bottom panel – lysates were treated with RNAse cocktail.

We validated the robustness of the assay by intracellular staining of cells expressing TurboID-nanobody, which revealed sufficient biotinylation even after 15 minutes exposure to biotin-supplemented medium (Fig. S4A). After intersecting the enriched proteins found in 3 biological replicates of the TurboID experiment, we narrowed the obtained list of proteins to 85 putative interactors (Fig. S4B), from which 81 were not previously associated with TENT5C in different systems^39,49^. Interestingly, even looking at all unique enriched proteins across 3 biological replicates, we could not find any peptides representing FNDC3A/B (Supplementary Data 2), which were previously established interactors of TENT5C in multiple myeloma^49^. Knowing that those proteins are expressed in erythroblasts, it was a strong indication that TENT5C operates differently in these cells. Notably, among enriched proteins, we found ∼40 to be associated with RNA binding Gene Ontology (GO) term (Fig. 4B). Among these, we found LARP4 and LARP5 to be the most specifically enriched after pulldown of biotinylated proteins (Fig. 4C). At the same time, PCBP1 and PCBP2, despite their known role in α-globin mRNA metabolism, were not enriched. LARPs are known to bind mRNAs and protect their poly(A) tails, making them intriguing candidates for future research, especially since their role in erythropoiesis has not yet been described. Among other interesting putative interactors, we found several potential regulators of TENT5C pointing to degradation of the protein by the ubiquitin-proteasome system (UPS), which is in line with previous findings and protein sequence composition of TENT5 family^49^ (Supplementary Data 2).

We validated the possibility of TENT5C and LARP4/5 interaction in the HEK293T system using ectopic overexpression of both 3xFLAG-TENT5C and HA-LARP4 or HA-LARP5 constructs followed by co-IP using anti-FLAG antibody. Notably, we found that LARP5 co-purifies with TENT5C in both RNAse-treated and untreated conditions (Fig. 4D). We observed that in the case of co-expression of 3xFLAG-TENT5C and HA-LARP4, there is a low protein abundance of both proteins visible in inputs, however, some weak signal corresponding to correct molecular weight can be seen for both HA-LARP4 and 3xFLAG-TENT5C in elution (Fig. 4D).

To sum up, we found novel putative interactors of TENT5C in erythroblasts, pointing to its ER-independent mechanism of action. This finding is in accord with cytoplasmic globin transcripts being the main substrates of TENT5C in erythroblasts. We concluded that LARP5 and TENT5C interaction is biochemically possible and independent of RNA presence.

### LARP4/5 cooperate with TENT5C and ameliorates its deficiency

To elucidate the potential role of LARP4/5 in erythropoiesis, we silenced them in WT erythroblasts with shRNA expressed from retroviral vectors (Fig. S5D). To better understand the potential cooperation of TENT5C and LARP4/5, we also used double depletions of either TENT5C and LARP4 or TENT5C and LARP5 and triple silencing of all genes. Since the cDNA sequencing of erythroblasts sorted from the BM revealed the strongest effect of TENT5C deficiency on poly(A) tails mostly in orthoE, we sorted CD71^low/-^, TER119^+^ populations of shRNA expressing erythroblasts cultured *ex vivo* for 48h.

Although silencing of LARPs in erythroblasts did not affect the major trajectory of differentiation (Fig. S5A), in the case of LARP5 and/or TENT5C depletion, we observed more CD71^low/-^, TER119^+^ cells compared to the control silencing (Fig. 5A), resembling the phenotype of *Tent5c^ko^ in vivo* (Fig. 1B). Such an effect was not seen for LARP4 depletion. This observation strongly suggests that faster erythropoiesis progression in *Tent5c^ko^* is triggered by intracellular factors rather than altered cytokine levels. Gratifyingly, looking at poly(A) tails with cDNA sequencing, we observed consistent poly(A) tail shortening across all conditions for combined globin mRNAs (Fig. 5B), supporting the role of LARP4/5 in poly(A) tail length regulation. This is in line with the observed decrease in abundance of globin mRNA reads in all conditions compared to the control (Fig. 5C, 5D). The most pronounced effect on poly(A) tails was observed for LARP5 depletion (Fig. 5B), accompanied by the downregulation of *Tent5c* mRNA (Fig. S5B). Accordingly, LARP5 depletion had the most pronounced effect on globin mRNA abundance visible in RT-qPCR (Fig. S5B). At the same time, even though LARP4 depletion caused weak shortening of the poly(A) tails (Fig. 5B), it had the opposite effect visible on qPCR with slight upregulation of TENT5C and globin mRNA abundance (Fig. S5B).

**Figure 5.**
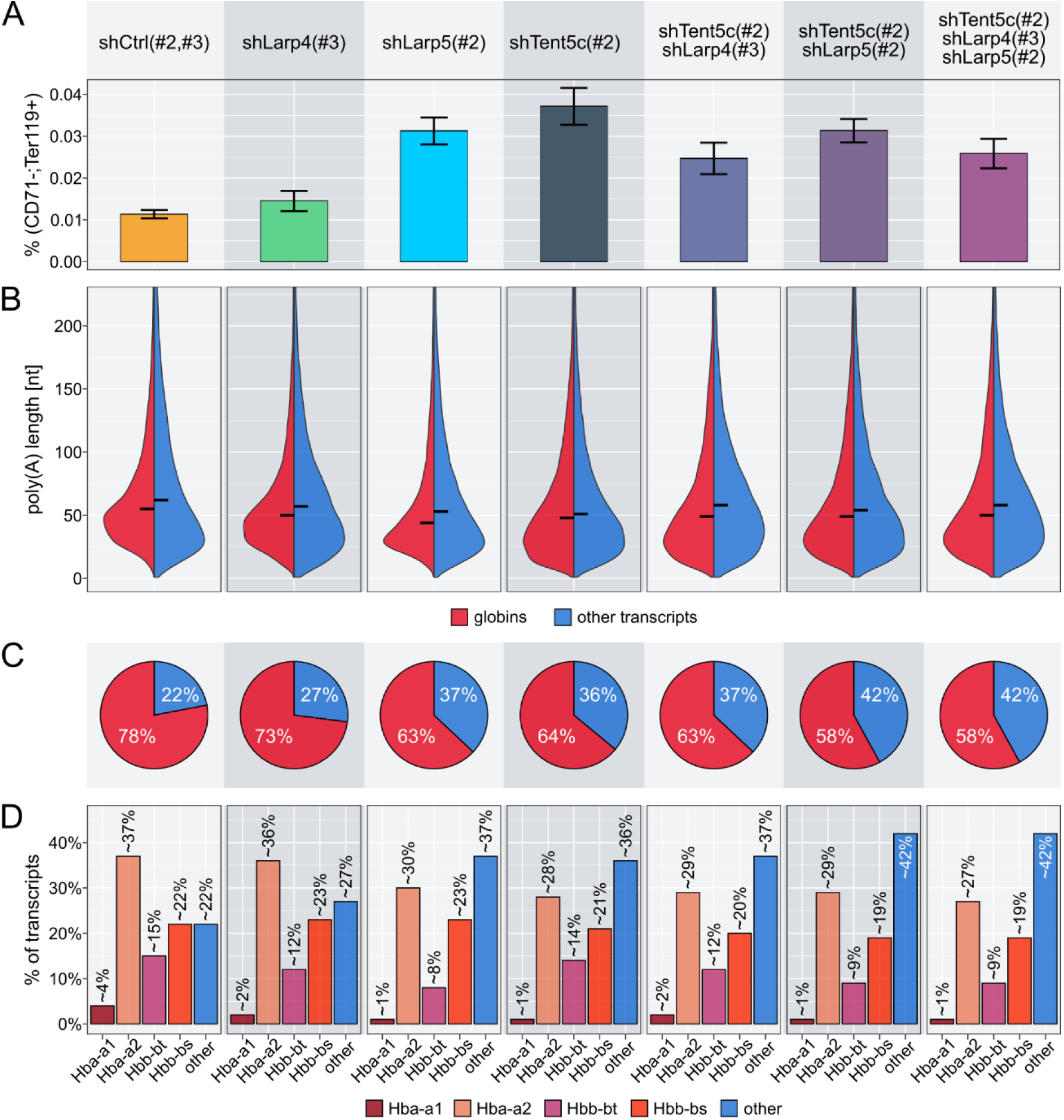
Functional association of LARP4/5 with globin mRNAs stability regulation. (A) Flow cytometry results summarizing the percentage of late erythroblasts (CD71-; Ter119+) in populations expressing selected shRNAs. (B) Distribution of poly(A) tail lengths in globin (red; left) and other (blue; right) transcripts in transfected cells. Black horizontal line indicates median. (C) Percentage of globins versus other transcripts in transfected cells. All globins are shown together versus the remaining transcripts. (D) Percentage of globins and other transcripts in transfected cells. Each hemoglobin is shown separately versus the remaining transcripts.

Despite the near complete depletion of TENT5C (Fig. S5C), we saw a mild effect of its silencing on globin transcript abundance (Fig. S5B). The effect size resembled that of *Tent5c^cat^* mutant erythroblasts cultured *ex vivo*, where we can observe upregulation of both LARPs, pointing to compensatory mechanisms (Fig. S5E).

Double depletion of TENT5C and LARP4 nullified the effect we observed in single LARP4 depletion (Fig. S5B). Similarly, the effect on poly(A) tails in double depletion of LARP4 and TENT5C for all globins combined was additive (Fig. 5B).

In the case of LARP5 and TENT5C silencing, globin abundance was similar to a single depletion of LARP5 (Fig. S5B). Consistently, we could see that the globin poly(A) tails were not further shortened in double depletions of LARP5 and TENT5C nor triple depletions of LARP4, LARP5 and TENT5C compared to LARP5 knock-down (Fig. 5B).

We concluded that both TENT5C and LARP4/5 are important for the stability of globin mRNAs, with LARP5 having a dominant effect. Given that LARP5 silencing causes co-depletion of TENT5C mRNA, we cannot draw a definitive conclusion regarding their mechanistic dependency. However, our results suggest that cytoplasmic polyadenylation driven by TENT5C is dependent on LARP5 rather than LARP4, differentiating these highly similar proteins. We propose that LARP4/5 protect globin mRNAs from deadenylation, in agreement with the ability of this family of proteins to directly bind mRNA poly(A) tail 3’ end, while, as we have already shown, TENT5C re-adenylate globin mRNAs.

### TENT5C is highly unstable in erythroblasts

Since the determination of TENT5C and LARP5 dependency is challenging and their interaction may be transient, further exploration of TENT5C regulation is needed to better understand its mechanism of action and the nature of possible interactions. *Tent5c* mRNA in late erythroblasts is highly expressed, representing the most abundant mRNA encoding poly(A) polymerase^28^. At the same time, TENT5C protein levels seem surprisingly low as proteomics analyses of murine and human erythroblasts have failed to identify a single peptide of TENT5C^3,4^. Therefore, we reasoned that TENT5C might be short-lived in erythroblasts. To test this, we utilized cycloheximide (CHX) chase assay on *ex vivo* culture of *Tent5c-EGFP* knock-in erythroblasts from the fetal livers. This experiment revealed a rapid downregulation of TENT5C-EGFP shortly after protein synthesis inhibition (Fig. 6A). Notably, 2.5h post-CHX treatment, the protein reached ∼10% of initial levels and was not depleted further (Fig. 6A).

**Figure 6.**
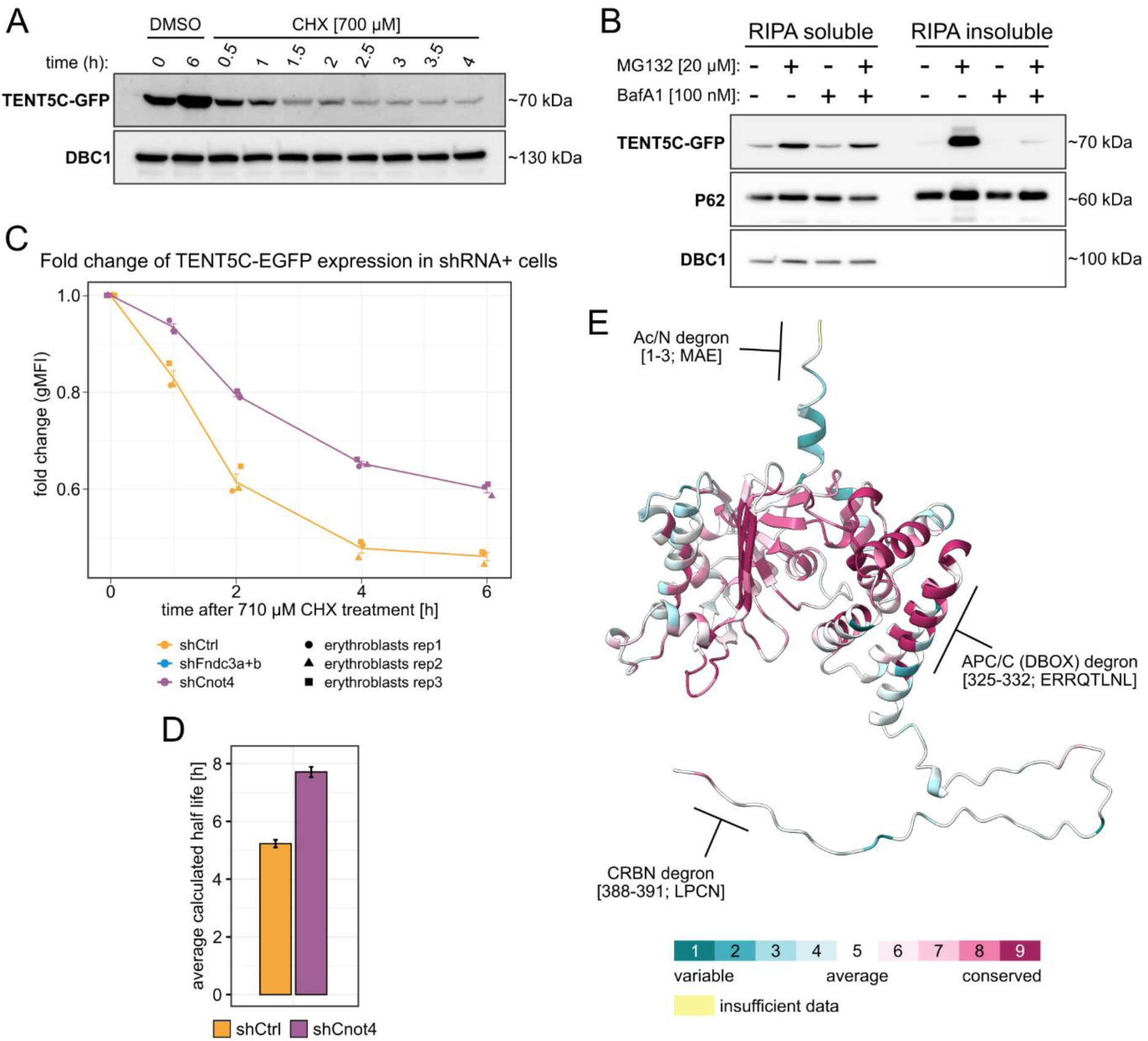
TENT5C instability and identification of contributing factors. (A) Western Blot showing a pulse-chase experiment upon cycloheximide treatment in erythroblasts. 2-leftmost samples represent DMSO treated cells gathered at the beginning and at the end of the experiment, remaining samples were gathered at denoted timepoints. (B) Western blot showing TENT5C-EGFP expression in erythroblasts upon inhibition of proteasomal degradation (MG132), autophagic degradation (BafA1) or both. The left side corresponds to RIPA-soluble fractions, right side corresponds to RIPA-insoluble fractions. (C) Line plots showing relative TENT5C-EGFP expression in subpopulations of cells expressing shRNA in pulse-chase cycloheximide experiment. Expression relative to initial levels (0h timepoint). Shape-coding applied for biological replicates, color-coding applied for condition. Data presented as mean ± SEM. (D) Bar plot representing calculated half-lives for each condition. Data presented as mean ± SEM. (E) Graphical representation of predicted degron motifs with exposed residues on TENT5C Alpha Fold 2 predicted structure. The structure is colored for evolutionary conservation score.

Our TurboID revealed several potential interactors which could be attributed to TENT5C regulation and/or degradation (p62, Anapc1, Otud4, Ubap2l, Ubap2, Bmp2k). Indeed, it has been demonstrated that TENT5C is degraded by the proteasome, and in the case of UPS inactivation, it is sequestered by p62, forming insoluble aggregates prone to autophagic degradation^49^. We observed similar behavior of TENT5C-EGFP in erythroblasts however, combining UPS and autophagy inhibition caused a weak effect on the accumulation within soluble fractions and unexpected loss of the TENT5C-EGFP protein in insoluble fractions (Fig. 6B).

Looking at the trigger of rapid TENT5C degradation in erythroblasts, we used a candidate approach. We focused on CNOT4 E3 ligase, a non-essential component of the CCR4-NOT deadenylating complex, which can be present close to mRNA poly(A) tails. The rationale was based on the identification of potential degron motifs within the TENT5C protein sequence, which revealed CNOT4 as a predicted candidate to act on the Ac/N degron found within the 3 N-terminal residues of TENT5C (Fig. 6E). Although, this motif was one of the least evolutionarily conserved among the predicted degrons with exposed residues, the consensus occurred in 88% of all mammalian orthologs aligned in the multiple-sequence alignment (MSA) (Supplementary Data 3). Notably, *Cnot4* silencing in erythroblasts increased the stability of TENT5C (Fig. 6C), with a calculated half-life exceeding 7 hours (Fig. 6D). Interestingly, degron motifs with the highest evolutionary conservation scores and exposed residues were also identified closer to the C-terminus (Fig. 6E), suggesting the possible involvement of multiple motifs in regulating TENT5C degradation.

In aggregate, we identified CNOT4 as one of the factors responsible for TENT5C instability in erythroblasts. Considering these results, we reason that the interaction of TENT5C with LARP5 could be restricted by TENT5C rapid degradation.

## DISCUSSION

This work, for the first time, describes the mechanism by which massive hemoglobin production is ensured until the last steps of terminal erythropoiesis. Notably, although the links between TENT5C and RBC development were previously suggested^46,51^, the molecular mechanism leading to the microcytic anemia phenotype upon TENT5C dysfunction remained elusive. Herein, we provide compelling evidence that this phenotype depends on the direct re-adenylation of globin mRNAs and, therefore, hemoglobin production rather than secondary defects like problems with iron uptake. Furthermore, we identify persistent stress erythropoiesis in the spleen of TENT5C-deficient mice as a compensatory mechanism triggered by insufficient hemoglobin production.

RBC development is an outstanding example where poly(A) tail homeostasis is crucial to coordinate the supply of mRNA and to allow its proper distribution throughout differentiation. The gradual shortening of the globin mRNA poly(A) tails during stages of erythropoiesis described by us correlates well with a known decrease in transcriptional activity^28^. Such correlation indicates that in reticulocytes where poly(A) tails are the shortest with 20A on average, globin mRNAs are prone to degradation through a deadenylation-dependent pathway (Fig. 3E, top panel). As seen for *Tent5c^cat^* reticulocytes there is a substantial decrease in globin transcripts abundance (Fig. 3E, bottom panel). Hence, we reason that TENT5C activity in earlier stages grants globin mRNAs enough stability to avoid premature degradation post-enucleation. After all, it is known that circulating reticulocytes are translationally active up to 2 days after being released to the bloodstream without the possibility of producing nascent transcripts^34^.

An interesting aspect of our studies is the difference between α– and β-globin mRNAs, which was also the subject of previous studies in the context of other trans-acting factors^37,52,53^. In the case of beta globins, the effect of TENT5C dysfunction on translation is already seen in polyE erythroblasts despite no visible change in their poly(A) tails or transcript abundance (Fig. S3A, 3F). This observation supports cytoplasmic polyadenylation as an important factor for efficient translation. On the other hand, the upregulation of α-globin protein levels in *Tent5c^cat^* persists from basoE erythroblasts where elongation of *Hba-a1* poly(A) tails occurs (Fig. S3A, 3F). This elongation, most likely resulting from transcriptional compensation, may account for increased alpha globin protein levels, highlighting the differences in mRNA stability mechanisms between globins^53^. Notably, α-globin mRNA stability greatly relies on the TENT5C-independent αCP mRNP complex, mutations in which components are associated with α-thalassemia^37^. In contrast, the 3’UTR of β-globin lacks structural elements to support the assembly of a complex comparable to the αCP. Instead, the two elements responsible for its stability are independent and redundant^52^. Interestingly, none of the etiological trans-acting factors for β-thalassemia has been directly linked to mRNA stabilization in the cytoplasm^54^. Our findings broaden the current knowledge on globin mRNA stability regulation and translation. Cytoplasmic polyadenylation driven by TENT5C is not only important to sustain hemoglobinization in reticulocytes, but it is also crucial for the efficient translation of β-globin, which is affected independently by its mRNA abundance. The observed molecular phenotype resembles β-thalassemia minor, which often manifests as a mild form of microcytic hypochromic anemia^55^, closely resembling that of *Tent5c^cat^* mice (Fig. 1A).

Looking at the mechanism of TENT5C-mediated polyadenylation through proteomic experiments followed by functional studies, we demonstrated that in erythroblast, TENT5C cooperates with LARP4/5, providing a new perspective into how TENT5C-driven mRNA stability regulation can proceed outside of the well-established ER-dependent mechanism shown in different model systems^39,42–45^. At the same time, cytoplasmic polyadenylation in erythropoiesis is very dynamic as TENT5C is highly unstable and operates through a transient interaction with LARP5. The instability of TENT5C can be attributed to CNOT4 E3 ubiquitin ligase, a subunit of the CCR4-NOT deadenylating complex, whose activity opposes polyadenylation^56,57^. Hence, we envision that deadenylation can supersede TENT5C-driven polyadenylation as the CCR4-NOT complex also could degrade TENT5C. On the other hand, the balance of poly(A)-dependent decay and stabilization can be shifted towards stabilization if the expression of TENT5C is high enough, as we observe in orthochromatic erythroblasts, where *Tent5c^cat^* cells exhibit the most pronounced effect for globin poly(A) tails (Fig. 3E, S3A, S3B). Ultimately, the fate of every mRNA is to be degraded; hence, in a system where the terminal stage of differentiation is marked by complete clearance of organelles and mRNAs, the beneficial effect of TENT5C may act only in a precisely defined time window.

The exact mechanism of the TENT5C and LARP4/5 cooperation is yet to be determined, especially since LARP4/5 are large, multi-domain proteins with long disordered regions and several folded domains known for their ability to bind RNA^58,59^. Notably, so far, LARPs have not been associated with cytoplasmic polyadenylation, and the poly(A) protective activity has been mainly studied in the case of LARP4^60^, where the semi-disordered N-terminal region is required for both PABPC interaction and poly(A) binding^61^. Interestingly, this region does not contain any conventional RNA-binding domain. Yet, it is a major determinant for poly(A) binding in contrast to the better-studied La module, which plays a minor role in this matter^61^. In contrast to LARP4, LARP5 has only been shown to promote translation through the stabilization of mRNAs with AU-rich 3’UTRs^62,63^. However, both LARPs share domains interacting with PABPC and a ribosome-associated kinase RACK-1, potentially linking the 3’ ends of mRNAs with initiating ribosomes^62^. As demonstrated in this work, the pronounced effect on globin poly(A) tails seen with LARP5 silencing could not be further enhanced by TENT5C depletion (Fig. 5B, S5B), indicating that LARP5 not only protects globin poly(A) tails but also facilitates TENT5C-driven polyadenylation, a novel function for LARP5.

In aggregate, we described a dedicated cytoplasmic polyadenylation mechanism underscoring the uniqueness of terminal erythropoiesis with respect to the regulation of mRNA stability and translation. Further research is needed to uncover a more detailed regulatory network of globin mRNA stability control, considering both deadenylation, re-adenylation, and poly(A) protection, as well as trans-acting factors dictating the stability of elements regulating poly(A) homeostasis.

## STAR METHODS

### Mice

The following mice strains were used in this study: B6CBAF1;B6-TENT5C KO/Tar; B6CBAF1;B6-TENT5C Cat (D90N;D92N)/Tar; B6CBAF1;B6-TENT5C-EGFP/GFP/Tar (Key Resource Table). Mice strains were generated and genotyped by Genome Engineering Unit, IIMCB (www.geu.iimcb.gov.pl) developed from Mouse Genome Engineering Facility (www.crisprmice.eu). All mice were bred in the animal house of Faculty of Biology, University of Warsaw. Mice were maintained in conventional conditions in open polypropylene cages filled with wood chip bedding (Rettenmaier). Environment was enriched with nest material and paper tubes. Mice were fed at libitum with standard laboratory diet (Labofeed B, Morawski). Humidity in the rooms was kept at 55 ± 10%, temperature at 22 °C ± 2 °C, at least 15 air changes per hour, and light regime set at 12 h/12 h (lights on from 6:00 to 18:00)^42^. Mice of both sex were sacrificed at age 12-20 weeks to isolate bone marrow and spleens, to perform experiments on fetal livers, pregnant females were sacrificed at embryonic development days E12.5-E15.5. All procedures were approved by the I Local Ethical Committee in Warsaw affiliated at University of Warsaw, Faculty of Biology (approval number WAW/693/2018), with the requirements of the EU (Directive 2010/63/EU) and Polish (Act number 266/15.01.2015) legislation.

### Tissues collection and blood analysis

Blood samples taken from facial vein were collected from 12-20-weeks old WT and *Tent5c^cat^* mice of both sexes and were commercially analyzed at Veterinary Diagnostic Laboratory LabWet in Warsaw (http://www.labwet.pl) on the day of blood collection. For flow cytometry analysis of circulating reticulocytes. In parallel serum samples to analyze cytokines levels were collected to Microvette 500 CAT-Gel (cat# 20.1344, Sarstedt). All mice were sacrificed by cervical dislocation. Adult livers, fetal livers, spleens, and femur and tibia bones were isolated immediately and kept on ice until further processing. Fetal livers were disrupted by trituration in cold PBS followed by centrifugation (800 × g /5 min/4°C), 10% of the sample was used for Flow Cytometry staining and the rest was snap frozen as pellets until further processing. Spleens were pressed by 70 µm cell strainer (352350, Corning) to obtain homogenous suspensions. Bone marrow was isolated by centrifugation (14000 × g / 30 s/ RT) of the isolated bone with cut-off tip of the bone head facing down the bottom of a smaller tube with a pinhole (0,5 ml) inserted into a 1,5 ml tube^39^.

EBIs from bone marrow and spleens were isolated as follows. Bone marrow was flushed from femurs, cells were harvested in Iscove’s Modified Dulbecco’s Medium (IMDM) (21980065, Gibco) medium supplemented with 20% fetal bovine serum. Cells aggregates were gently pipetted using 1 ml pipet and filtered through a 70 µm cell strainer. Spleens were minced and cell suspension was filtered through a 70 µm cell strainer. Cells suspensions from both bone marrow and spleens were fixed with 4% paraformaldehyde in PBS for 10 min. Cells were washed with FACS buffer (PBS supplemented with 0.2% BSA) and stained with antibody mix against EBIs described in flow cytometry section.

### Flow cytometry

Unless stated otherwise, cells (1.5 × 10^6^) were incubated with Fc Block (anti-CD16/32) for 10 min RT followed by washing in FACS buffer (0.2% BSA in PBS) with centrifugation (800 × g / 5 min / 4°C). All antibodies used are listed in key resource table. All staining mixes were prepared in 50 µl brilliant Stain Buffer (563794, BD Biosciences). Cells were stained by incubation with staining mix for 40 min in 4°C protected from light followed by washing in FACS buffer. Next, cells were stained with a chosen LIVE/DEAD fixable viability dye (Invitrogen) specified for each cytometry panel according to manufacturer’s recommendation. Cells were washed in FACS buffer and resuspended in 500 µl FACS buffer. Samples were measured using CytoFLEX LX (Beckman Coulter Life Sciences) under CytExpert v2.2 software control and sorted using CytoFLEX SRT (Beckman Coulter Life Sciences) under CytExpert v1.1 software control.

Cells isolated from fetal liver (Fig. S1A, S1D, 3A), were stained with anti-CD3 BV421, anti-CD11b BV421, anti-CD41 BV421, anti-CD45R BV421, anti-Ly6G/Ly6C BV421, anti-CD71 PE, anti-Ter119 Alexa Fluor 647 antibodies according to supplier’s recommended antibody concentration. To discriminate dead cells LIVE/DEAD Fixable Near-IR Dead Cell Stain (L34994, Invitrogen) was used. The gating strategy of cells isolated from fetal liver and *ex vivo* cultures is presented in Fig. S7A^12,13^.

Cells isolated from bone marrow and spleens (Fig. 1B, S1C, 2F, 3B) were stained using the same antibody mix and Live/Dead stain as in case of fetal liver but with addition of anti-CD44 PE-CF594 antibody. Gating strategy for adult erythropoiesis is presented in Fig. S7B^5,12^. To quantify the expression of TENT5C-EGFP both gating strategies were used (Fig. S7 gating A, B). TENT5C-EGFP expression was calculated as gMFI of cells expressing GFP diminished by autofluorescence of WT cells.

Cells from blood were stained with anti-CD71 FITC antibody to estimate the amount of circulating CD71^+^ immature reticulocytes in the blood stream (Fig. 1C).

To analyze EBIs (Fig. 2E, 2G, 2H) the following antibodies were used for staining: anti-CD11b BV786, anti-F4/80 BUV395, anti-Ly6C PE-Cy7, anti-VCAM1 BV650, anti-CD169 BV421, anti-CD71 PE, anti-Ter119-Alexa Fluor 647. To discriminate dead cells LIVE/DEAD Fixable Near-IR Dead Cell Stain kit was used (L34994, Invitrogen). Gating strategy for EBIs calculation assumes that true EBIs are cellular aggregates positive for both erythroblastic and macrophages markers^9,10,64,65^, and is shown in Fig. S7C. Gating strategy for characterization of single macrophages is also shown in Fig. S7C. Bone marrow and spleen EBIs analysis was performed by calculation of percentages of CD71^+^Ter119^+^F4/80^+^VCAM1^+^ within the live gate^10,64^.

Erythroblasts from *ex vivo* culture expressing shRNA against *Tent5c* and *Larp4/5* (Fig. 5, S5A) were stained with: anti-CD71 FITC, anti-Ter119 Alexa Fluor 647 and LIVE/DEAD Fixable Violet Dead Cell Stain Kit (L34964, Invitrogen). To sort and analyze late erythroblasts from *ex vivo* culture expressing shRNA we used gating strategy shown in Fig. S7E.

For flow cytometry based CHX-chase assay (Fig. 6C) and to test shRNA depletion efficiencies (Fig. S5C, S5D, S6C) cells from *ex vivo* culture were stained only with LIVE/DEAD Fixable Violet Dead Cell Stain Kit (L34964, Invitrogen) without blocking. To calculate TENT5C-EGFP expression during silencing of *Tent5c/Fndc3a+Fndc3b/Cnot4* with shRNA in *ex vivo* cultured cells, we used gating strategy shown in Fig. S7D. To test depletion efficiency by western blotting cells expressing shRNA were sorted according to the same gating strategy.

To evaluate the robustness of the TurboID assay (Fig. S4A), we fixed 5 × 10^5^ of acquired cells and stained them with Streptavidin, AlexaFluor-647 Conjugate (S21374, Invitrogen) at 10 µg/µl working concentration using the reagents and following the guidelines of BD Cytofix/Cytoperm Fixation/Permeabilization Kit (554714, BD Biosciences).

To quantify cytokine levels in *Tent5c^ko^* (Fig. 2A), we used commercially available cytometric bead arrays: LEGENDplex Mouse HSC Erythroid Panel (740681, BioLegend), LEGENDplex Mouse Cytokine Panel 2 (740134, BioLegend), LEGENDplex Mouse Anti-Virus Response Panel (740621, BioLegend).

### Measurement of iron levels

Serum iron levels (Fig. S1B) were measured with the SFBC kit (80008, Biolabo) according to the manufacturer’s protocols. To measure tissue non-heme iron content, the bathophenanthroline method was used^66^. Briefly, tissue samples were dried at 45°C for 72 hours, and then digested in a 10% HCl/10% TCA acid solution at 65°C for 48 hours. Iron concentration in the solution was determined by measuring absorbance at 535 nm after reaction with the chromogenic reagent (6M sodium acetate, 0.01% bathophenanthroline-disulfonic acid, 0.1% thioglycolic acid) and calculated relative to tissue dry weight using a standard curve.

### Cell lines

HEK293T (CRL-3216, ATCC) cells were cultured in DMEM high glucose medium (41965039, Gibco) supplemented with 10% fetal bovine serum (FBS) (10270-106, Gibco). In the case of HEK293T cultured for virus production the medium was additionally supplemented with 1% Penicillin/Streptomycin (P0781, Sigma). Cells were cultured in incubator set to 37°C, 5% CO2. Cells were passed after reaching 90% confluence by incubation with Trypsin / EDTA 2 (C-41012, Sigma-Aldrich) for 5min at 37°C.

### Primary cell culture of murine erythroblasts

Erythroid progenitors from E14.5 fetal livers were isolated using Ter119 negative selection, with random pooling of the livers. Progenitors obtained from a separate pool were considered a biological replicate. Up to 6 fetal livers were pooled in 10% FBS/PBS solution and disrupted by pipetting to obtain homogenous cell suspension, followed by straining through 70 µm nylon strainer (352350, Corning). Cells were kept on ice throughout isolation, and always washed or incubated with 10% FBS/PBS solution (wash buffer). After centrifugation (800 × g /5 min/4°C), cells were resuspended in 1ml wash buffer supplemented with 10 µl Rat IgG [10 µg/µl] (I4131, Sigma) and incubated on ice for 15 min. Without washing, 2µl of biotinylated anti-TER119 antibody [0,5 µg/µl] (13-5921-82, Invitrogen) per 1 fetal liver was added to the cell suspension and incubated for 15 min on ice with gentle agitation from time to time. Cells were washed with 10 ml wash buffer and resuspended in 1ml wash buffer. Dynabeads MyOne Streptavidin C1 (65001, Invitrogen) were washed twice with 10% FBS/PBS before incubation with cells and resuspended in original volume with wash buffer. 30 µl Dynabeads MyOne Streptavidin C1 was used per 1 µl of anti-TER119 antibody. Cells were incubated with beads for 10 minutes on ice. Next, cell suspension was brought to final volume of 4 ml and separated on EasyEights EasySep Magnet (18103, STEMCELL Technologies). Supernatants were subjected to another round of separation to avoid carryover of residual magnetic beads. TER119^+^ fractions bound to magnetic beads were retained and frozen. Supernatants containing TER119^-^ cells were centrifuged and pellets resuspended with 10ml wash buffer for counting. Cells were counted using Countess 3 FL Automated Cell Counter (AMQAF2000, Invitrogen) with Trypan blue stain (T10282, Invitrogen) to assess viability. Following final centrifugation, supernatant was rejected and cells were resuspended in erythroblasts differentiation medium based on IMDM (21980065, Gibco) containing 2 U/ml Recombinant Human EPO (100-64, Peprotech), 20% FBS (10270-106, Gibco), [10 µg/ml] insulin (I3536, Sigma), 200 µg/ml holo-transferrin (T0665, Sigma), 100 µM 2-Mercaptoethanol (M3148, Sigma) and 1% Penicillin/Streptomycin (P0781, Sigma) with seeding density of 4 × 10^5^ cells/ml^16^. Cells were cultured in untreated 12-well plates (CLS3737-100EA, Corning) in incubator set to 37°C, 5% CO2.

### Primary cell culture of B-cells

B-cells were isolated using EasySep Mouse B Cell Isolation Kit (#19854, STEMCELL Technologies). Spleens from adult mice were collected on ice and pressed against nylon strainer in PBS to create cell suspension. Cell suspension was centrifuged (800 × g /5 min/RT) and the remaining procedure was carried out as described in the manufacturers protocol. Isolated B-cells were counted and seeded with starting density of 1M live cells / 1ml medium. Cells were cultured in RPMI 1640 ATCC modification (#A1049101, Invitrogen) with 15% FBS (10270-106, Gibco), 1% penicilin/streptomycin (P0781, Sigma), 100 µM 2-Mercaptoethanol (M3148, Sigma) and 20 µg/ml LPS (#L2630, Sigma). Culture medium was changed every day and culture density was maintained at 1M live cells / 1ml medium through the entire culture. On day 2 of culture cells were infected with retrovirus expressing shRNA as described in separate section. 2 days post infection shRNA expressing cells were analyzed and sorted with FACS.

### Molecular cloning

All vectors used in the work were generated by SLIC and/or PCR cloning. In every case we used NEB Stable Competent E. coli (C3040I, New England Biolabs) for transformation. In the case of shRNA cloning, we used T4 DNA ligase (EL0011, Thermo Scientific) to ligate annealed and phosphorylated oligos to dephosphorylated linearised pMSCV-mCherry-BbsI-U6 vector. All plasmids and oligos encoding shRNA are listed in key resources table. pMSCV-EGFP-BbsI-U6 vector which served as a backbone to create pMSCV-mCherry-BbsI-U6 was a gift from Prof. Harvey Lodish.

### Plasmid isolation

Plasmids were propagated in NEB Stable Competent E. coli (C3040I, New England Biolabs). Plasmids for transfection were isolated with Plasmid Midi AX kit (092-10, A&A Biotechnology) according to manufacturer’s protocol. Plasmids for cloning were propagated using smaller format Plasmid mini (020-250, A&A Biotechnology).

### HEK293T transient transfection for co-IP

Cells were seeded on 6-well plates (0.5*10^6^ cells per well) in 2 ml of DMEM medium with 10% FBS for 20-24h before transfection. For immediate induction of the transgenes, the medium was supplemented with Tetracycline Hydrochloride (A39246, Gibco) 2 hours before transfection creating working concentration of 100 ng/ml. Transfection was made with TransIT-2020 Transfection Reagent (MIR5400, Mirus) reagent. Briefly, 250 µl of OptiMEM (31985070, Gibco) was mixed with 1,3 µg of pKK plasmid encoding GOI. Then 2 µl of transfection reagent was added and the mixture was incubated for 20 min at room temperature, followed by a dropwise delivery to the cells. Cells were collected for experiments 48h post-transfection.

### Co-immunoprecipitation

Co-immunoprecipitation (co-IP) of FLAG-tagged proteins was performed with ChromoTek DYKDDDDK Fab-Trap Agarose (ffa, Proteintech) and ChromoTek Binding Control Agarose Beads (bab, Proteintech) as a binding control (BC).

HEK293T cells (3 wells in a 6-well plate per condition) were collected 48h after transfection and suspended in 1 ml Lysis Buffer (50 mM Tris-HCl pH 7.5, 150 mM NaCl, 0.5% NP-40, 0.5 mM EDTA) supplemented with 1 mM PMSF, 20 nM pepstatin, 6 nM leupeptin, 2 ng/ml chymostatin. Then, the material was divided into two tubes and to one of them, 20 µl of RNase Cocktail Enzyme Mix (AM2286, Ambion) was added. Samples were incubated for 30 min at 4°C on the rotator. Lysates were centrifuged (10000 × g/15 min/4°C) and supernatant was used for immunoprecipitation. 25 µl of resin per sample was washed 4x with Lysis Buffer and then incubated with cell lysate for 1.5 h at 4°C on the rotator. Then, samples were centrifuged (2500 × g/5 min/4°C), supernatant was collected as a flow-through fraction, and resin was washed 3 times with lysis buffer. During the third wash, samples were transferred to a new tube. For elution, the resin was suspended in 40 µl of 2x Laemmli Buffer (0.125 M Tris base, 0.14 M SDS, 20% glycerol, 10% 2-mercaptoethanol, 0.002% Bromphenol blue), denatured for 10 min at 100°C, centrifuged and the supernatant transferred to a new tube.

### Protein extraction and western blotting

Cells were harvested on ice and washed twice with cold PBS. To lyse 1 × 10^6^ cells we used 100 µl RIPA buffer (150 mM NaCl, 1% v/v Triton X 100, 0.5% sodium deoxycholate, 0.1% SDS, 50mM Tris pH 8, 0,5 mM EDTA, pH = 8) supplemented with protease inhibitors: 1 mM PMSF, 20 nM pepstatin, 6 nM leupeptin, 2 ng/ml chymostatin. Cells acquired by FACS were sorted straight to 4x concentrated RIPA buffer (1 × 10^4^ cells / 40 µl buffer) supplemented with 4x concentrated protease inhibitors. Lysis was performed on ice for 15 min with gentle mixing from time-to-time following clarification by centrifugation at 15 min/4°C/14000 × g. Supernatant (RIPA soluble fraction) was retained, and protein concentration was measured with Pierce BCA Protein Assay Kit (23225, Thermo Scientific), according to manufacturer’s protocol. The remaining pellet was either discarded or washed twice with RIPA buffer followed by another round of solubilization in 50 µl of 5% CHAPS in 7M urea, supplemented with the same inhibitors as described above. Pellets were kept on ice for 30 min with gentle agitation from time to time. Afterwards, lysates were clarified by centrifugation 15 min/4°C/14000 × g and supernatants were retained (RIPA insoluble fraction). RIPA insoluble fractions were not routinely quantified for protein as they had too low concentration. In the case of RIPA soluble fractions samples were adjusted to the same concentration and heated for 7 min 98°C with appropriate volume of 4x Laemmli sample buffer (0.25 M Tris base, 0.28 M SDS, 40% glycerol, 20% 2-Mercaptoethanol, 0.004% Bromophenol blue). RIPA insoluble fractions were treated as if they had the same protein concentration, otherwise they were processed the same. An equal sample volume was applied on denaturing SDS gel. Electrophoresis was performed until desired gel resolution was obtained. The protein was transferred onto the 0.2 µm nitrocellulose membranes (10600001, Amersham) using wet transfer method (25 mM Tris, 192 mM glycine, 20% methanol, 0.2% SDS) with parameters set to constant 400 mA for 1 h 15 min at 4°C. After transfer, the membranes were stained with Ponceau stain (Ponceau S 0,1 % (w/v) in 5% acetic acid) for 5 min at RT, followed by washing in ddH_2_O and staining was documented. Membranes were blocked in 5% skimmed milk / TBST (w/v) for 1h at RT. Membranes were incubated with primary antibodies with addition of 0.05% sodium azide, rotating overnight at 4°C. All primary antibodies used for WB in the study (listed in key resources table) were used at 1:1000 concentration except for anti-GAPDH which was used at 1:5000 concentration. Afterwards, membranes were washed 3 times in TBST (5 minutes each wash) and incubated for 1 h at RT with secondary antibodies (listed in key resources table) diluted 1:10000 in 5% skimmed milk / TBST followed by 3 washes in TBST. Membranes were developed with either Clarity Western ECL Substrate (#1705061, Bio-rad) or SuperSignal West Femto Maximum Sensitivity Substrate (34094, Thermo Scientific) if the signal was too weak to see the bands with standard substrate. Visualisation was done on either ChemiDoc MP Imaging System (Bio-rad) or iBright FL1500 Imaging System (Thermo Scientific) and the images were processed with appropriate manufacturer’s software.

### RNA isolation, purification and quality control

Frozen fetal livers were immediately dissolved in 900 µl TRIzol Reagent (15596018, Invitrogen) heated to 60°C by pipetting and vortexing until fully solubilized. Further steps of the isolation were carried out as described in manufacturers protocol. Chemical purity of RNA was determined with NanoDrop OneC (ND-ONEC-W, Thermo Scientific). Absorbance ratio A260/230 in range 1.9-2.3 was considered a pure sample. In case of impurities samples were subjected to purification on KAPA Pure Beads (07983298001, Roche) according to manufacturer protocol with no size-exclusion cut-off. Sample integrity was determined using High Sensitivity RNA ScreenTape (5067-5579, Agilent) on 2200 TapeStation system (G2964AA, Agilent). To isolate total RNA from cells acquired by FACS we used PicoPure RNA Isolation Kit (KIT0204, Applied Biosystems) with some modifications. 10000 cells were sorted directly into 150 µl of extraction buffer with final volume of 200 µl. Samples were mixed with 70% Ethanol in 1:1 ratio. The rest of the steps were carried out as described in the manufacturers protocol including DNAse treatment with DNAse I (79254, Qiagen). Samples were eluted in 15 µl elution buffer. 10% of the eluate was used to assess RNA integrity as described above; 20% was used to synthesize cDNA for RT-qPCR; 70% was mixed with 50 ng of chemically pure total RNA from *C. elegans* following purification on KAPA beads. The eluate was used for Nanopore cDNA library preparation.

### 5’CAP mRNA enrichment

To maximize the amount of sequencing material for DRS we used mRNA enrichment using the in-house purified mutant of eIF4E^K119A^ with increased CAP affinity^67,68^ tagged with a GST tag. All centrifugation steps were carried out in RT for 5 min at 500x g, unless stated otherwise. We used 250 ul Gluthatione-Sepharose 4B (17-0756-01, GE Healthcare) resin per enrichment of 50 µg of total RNA from the fetal liver. Resin was equilibrated by washing 2x in 1ml PBS. Supernatant was discarded and resin was resuspended in 200ul PBS. To resuspended resin 0,2 mg of mutant eIF4E was added to create final concentration of 1 mg eIF4E / 1 ml resin, following incubation with gentle rotation for 2h in RT. Next, resin was centrifuged and supernatant discarded, following 2 washes with 1ml PBS. Resin was washed 3x in 1ml buffer A (10 mM potassium phosphate buffer, pH 8.0 (PPB), 100 mM KCl, 2mM EDTA, 5% glicerol, 0.005% Triton X-100, 6 mM DTT, 20 U/ml Ribolock RNAse Inhibitor). Total RNA was heated to 70°C for 10 following cooling on ice for 2 min. The samples were filled up to 500 µl with buffer A and centrifuged resin was resuspended with sample solution. Samples were incubated with gentle rotation for 1h in RT. Next, samples were centrifuged, supernatant was rejected, and bound resin was washed 3 times with buffer A, following 2 washes with buffer A supplemented with 0.5 mM GDP. Samples, were resuspended in 1ml A and transfered to new tubes following an additional wash with buffer A. To elute mRNA centrifuged resin was resuspended in 400 µl buffer A, following addition of 500 µl phenol:chloroform:isoamyl (25:24:1) and vortexing for 10s. Samples were put on ice for 15 min, following centrifugation (17000 × g/10 min/4°C). Aqueous phase was transferred to a new tube and equal volume of chloroform was added, following mixing by inverting the tube 5 times. Samples were centrifuged 17000 × g/3 min/4°C. Aqueous phase was transferred to a new tube with 1ml 98% ethanol, 50 µl 3M sodium aceteate pH = and 1 µl GlycoBlue Coprecipitant (AM9515, Invitrogen), following vortexing. Samples were kept overnight in –20°C for precipitation. Next day, samples were centrifuged 17000 × g /30 min/4°C, following 2 washes with cold 70% ethanol with centrifugation 17000 × g /15 min/4°C. Pellet was air-dried under fume hood and 15 µl of RNAse free ddH_2_O was added. Samples were kept in –80°C, until further processed.

### RT-qPCR

1/3^rd^ of the DNAsed sample from sorted cells was used for reverse transcription using SuperScript III Reverse Transcriptase (18080093, Invitrogen) according to manufacturer’s protocol. After cDNA synthesis, sample was diluted 2,5x to final volume of 50 µl. We used 2 µl of diluted cDNA to quantify relative gene expression with qPCR using Platinum SYBR Green qPCR SuperMix-UDG (11733038, Invitrogen) in final reaction mix volume of 10 µl with primer working concentration kept at 250 nM. Reactions were pipetted on 384-well plates (4309849, Applied Biosystems) and ran on QuantStudio 5 Real-Time PCR System (A28140, Thermo Scientific) with thermal cycling program recommended in the manual of used reaction master-mix. Primers were designed to amplify a specific region spanning exon-exon junction using Primer-BLAST algorithm^69^. In the case of globin genes (*Hba-a1* and *Hba-a2* or *Hbb-bs* and *Hbb-bt)* the nucleotide sequence similarity between paralogs prevented us from discriminating the expression levels between them, hence generic primer pairs were used to estimate the expression of both paralogs of either α-or β-globin. All primers used for qPCR were tested for efficiency by serial dilutions of the matrix. Calculated efficiencies were near 100% and similar between primer pairs for which the differences never exceeded 5%. Specificity was determined based on dissociation curve analysis. Relative gene expression was calculated using ddCt algorithm^70^.

### TurboID proximity biotinylation assay

TurboID-nanobody construct was introduced to cells by lentiviral transduction. Virus production and transduction methods were described in separate sections. After 48h of culture, cells from 6 wells of 12-well plate were pooled (approximately 4 × 10^7^ cells) following washing with PBS and resuspension in 6ml erythroblasts differentiation medium (described in separate section) containing 500 µM biotin (B4501, Sigma-Aldrich) following 15 min incubation at 37°C. Cells were collected on ice to stop the biotinylation and all following steps including the lysate handling and pulldown were done in 4°C to avoid biotinylation *in vitro*.

Cells were washed with ice cold PBS and spun 800 × g/ 4°C for 5 min. Washing was repeated 2 times. To evaluate the robustness of the assay we fixed 5 × 10^5^ of acquired cells and stained them with Streptavidin fluorescent conjugate as described in flow cytometry section. The remaining cells were lysed in 1 ml RIPA buffer with protease inhibitors and protein concentration was quantified as described in the protein extraction section. All washes performed downstream were carried out by resuspension of the beads in specific buffer, 2min rotation and magnetic separation followed by rejection of the supernatant. For pull-down of biotinylated proteins we used Pierce Streptavidin Magnetic Beads (88817, Thermo Scientific) which were equilibrated by washing twice in RIPA buffer prior to incubation with the lysate. We used 1mg of equilibrated beads per 0,5 mg of protein with lysate final concentration kept at 1 mg/ml protein. Beads were incubated with the lysate overnight in 4°C. Beads were separated and supernatant was retained as flow-through fraction. Beads were subjected to a series of washes: 2x with RIPA devoid of Triton X-100, 1x in KCL, 1x in 100 mM Na_2_CO_3_, 1x in 2M urea, 2x with RIPA devoid of Triton X-100 and 2x in 50mM Tris-HCl pH = 7,5. After last wash beads were separated, supernatant was discarded following short centrifugation 10s / 10000 × g /4 °C and residual liquid was removed while the tube was on the magnetic stand. Dry beads were kept in –80°C until further processed.

### Sample processing and LC-MS/MS analysis

For each sample, 100 µL of 100 mM ammonium bicarbonate and 2.5 µL of 200 mM tris(2-carboxyethyl)phosphine (TCEP) were added. Samples were vortexed briefly and placed on a horizontal shaker set to 10,000 rpm at room temperature for 30 min. Next, 2 µL of methyl methanethiosulphonate (MMTS) were added, and the samples were shaken for an additional 20 min at room temperature. Trypsin/LysC mix, prepared in 8 M urea with 100 mM ammonium bicarbonate to a final enzyme concentration of 0.02 µg/µL, was added to the samples (50 µL per sample). The samples were incubated at 37°C for 4 h with shaking, after which 300 µL of 100 mM ammonium bicarbonate was added. Digestion was allowed to proceed overnight. Following digestion, the samples were acidified with 10 µL of 5% trifluoroacetic acid (TFA). Tryptic peptides were purified using a modified SP3 protocol^71^ and subsequently vacuum-dried. Samples were analyzed on an EvosepOne (Evosep Biosystems) online LC-MS system coupled to an Orbitrap Exploris 480 mass spectrometer (Thermo Fisher Scientific). Peptide mixtures were loaded onto C18 Evotip trap columns, following the manufacturer’s protocol. This involved activation of the sorbent with 0.1% formic acid (FA) in acetonitrile, a 2-minute incubation in 1-propanol, and equilibration of the chromatographic sorbent with 0.1% FA in water. Samples were loaded in 30 µL of 0.1% FA, with centrifugation at 600 × g for 1 minute after each step.

Chromatographic separation was performed at a flow rate of 500 nL/min using the 44-minute (30 samples per day) gradient on an EV1106 analytical column (Dr. Maisch C18 AQ, 1.9 µm beads, 150 µm ID, 15 cm long, Evosep Biosystems, Odense, Denmark). Data acquisition was carried out in positive ion mode using a data-dependent acquisition (DDA) method. MS1 scans were acquired at a resolution of 60,000 with a normalized AGC target of 300%, auto maximum injection time, and a scan range of 350–1400 m/z. MS2 scans were performed at a resolution of 15,000, using a top-40 method with a precursor isolation window of 1.6 m/z. Dynamic exclusion was set to 20 s with a mass tolerance of ±10 ppm, and the precursor intensity threshold was set to 5 × 1000. Fragmentation was achieved using higher-energy collisional dissociation (HCD) at a normalized collision energy of 30%. The spray voltage was set to 2.1 kV, with a funnel RF level of 40 and a heated capillary temperature of 275°C.

### shRNA design

shRNA was designed using Block-iT RNAi designer tool (Thermo Scientific). We chose 3 sequences predicted to have the best depletion efficiency. After initial optimalisation and checking for secondary effects by flow cytometry we chose 1 sequence with the best depletion efficiency that seemed to have no secondary effects with respect to differentiation trajectory of erythroid cells.

### Virus production in HEK293T

HEK293T cells maintained as described in cell lines section. Confluent 293T cells were passed 1:6 on p100 plates. Next day in the late evening cells were transfected by calcium phosphate method, where transfection mix containing 250 mM CaCl_2,_ packaging and transfer plasmids is added to 2xHBS (280 mM NaCl, 50 mM Hepes, 1.5 mM Na₂HPO₄, 10 mM KCl, 12 mM Sucrose, pH 7.5) at 1:1 ratio. Plasmids for lentiviral production in transfection mix were concentrated as follows: 30 ng/µl psPAX2, 10 ng/µl pMD2.G and 40 ng/µl transfer. Plasmids for retroviral production in transfection mix were concentrated as follows: 20 ng/µl pCL-Eco and 40 ng/µl transfer. In the case of double or triple depletions, equimolar amounts of transfer plasmids were combined in the transfection mixes. The transfection mix was added dropwise directly to 2xHBS while vortexing and further vortexed for another 10s after the mix was added. The solution was incubated for 15 minutes at RT. Next, the solution was mixed bv tirturation and 1ml was added dropwise per 1 plate of 293T cells, following mixing of the plate by hand. From this point cells were handled according to BSL-2 standard. Next day in the early morning the culture medium was discarded and changed to 5,5 ml fresh medium. Next day on the afternoon, the viral supernatant was harvested and ready for transduction of cells of interest.

### Viral transductions

shRNA and TurboID constructs were introduced to cells by lenti-or retro-virus transduction respectively. Virus production was described in the separate section. Erythroid progenitors / B-cells isolated according to abovementioned sections were resuspended in concentrated viral supernatant at density 800k cells / 10ml supernatant in the presence of 10 µg/ml polybrene (sc-134220, Santa Cruz Biotechnology) for erythroid progenitors or 6µg/ml for B-cells. Cell suspensions were centrifuged (800 × g / 1,5 h / 34°C). Supernatant was aspirated, and cells were resuspended in appropriate volume of culture medium as described in primary cell culture sections. Optimal virus supernatant volume: cell number ratio was determined by serial dilution of cells taken for transduction starting from 1 × 10^6^ cells. Cells expressing shRNA were selected by the presence of mCherry fluorescent reporter using FACS.

### Inhibitor treatments

All drugs used in this study were solubilized in DMSO. In every case control cells were incubated with the appropriate volume of vehicle, which was never higher than 0,5% (v/v).

In the case of MG132 (S2619, Selleckchem) and Bafa1 (tlrl-baf1, InvivoGen) treatments we used [20 µM] and [100 nM] concentrations respectively for 5h between 43 and 48 h of *in vitro* differentiation of erythroid progenitors. In the case of B-cells we used the same treatment time and concentrations of the drugs on day 4 of culture.

In cycloheximide (CHX) chase experiments, the working concentration of the drug was 710 µM (C1988, Sigma-Aldrich). Control treatments with DMSO was carried out only for the first (0h) and last time-points. For chase experiments shown in Fig. 6A erythroid cells were incubated with CHX between 44 h and 48 h of *in vitro* differentiation and collected in 0,5h increments. In the case of B-cells (Fig. 6B) treatment with CHX was done on day 4 post-activation and collected at 0h; 2h; 4h; 6h timepoints. Cells were collected and protein extracted according to the previous section. To evaluate the effect of various depletions on TENT5C stability (Fig. 6C) both cell types were collected at 0h; 1h; 2h; 4h; 6h post CHX treatment. Transduced erythroblasts were treated with CHX between 44h and 50h of *in vitro* differentiation post-transduction and B-cells were treated with CHX 48h post-transduction (day 4 of *in vitro* culture). Collected cells were immediately put on ice and washed with cold FACS buffer, following staining as described in the flow cytometry section.

### Nanopore DRS library preparation

Libraries for nanopore direct RNA sequencing (DRS) were prepared using SQK-RNA002 Direct RNA sequencing kit (ONT) according to manufacturer’s instructions. We used 2 µg of CAP-enriched mRNA acquired by method described above. To improve sequencing performance and efficiency 150–250 ng of *Saccharomyces cerevisiae* oligo(dT)-enriched mRNA was added to all samples. To control the sequencing quality and throughput, the samples were spiked with 25 ng of synthetic oligonucleotide construct (instead of control strand supplied with kit). Sequencing experiments were run on the flow cell FLO-MIN106D (R9.4.1) using MinION (ONT) device controlled with MinKNOW (ONT) software. Obtained data were then basecalled with Guppy (ONT).

### Nanopore cDNA library preparation

Libraries for nanopore cDNA sequencing were prepared using SQK-PCB111-24 barcode cDNA sequencing kit (ONT) following the manufacturer’s protocol. Sequencing was performed on the flow cell FLO-MIN114 (R10) with the MinION (ONT) device controlled by MinKNOW (ONT) software. Obtained reads were basecalled using Dorado 0.7.0 (ONT).

### Data analysis

Statistical and exploratory data analysis of the results related to Fig. 1; Fig. S1; Fig. 2; Fig. 3A, 3B was performed using GraphPad Prism 6 software. Unless stated otherwise, the remaining data analysis was performed in R environment with details described below.

### Nanopore data analysis

Basecalled DRS reads were mapped to the reference transcriptome (Gencode vM32) using minimap2^72^ with following parameters: –k 14 –ax map-ont –-secondary = no. The resulting files were then processed with samtools^73^ to filter out supplementary alignments and reads mapping to the reverse strand (samtools view –b –F 2320). The lengths of the poly(A) tails were assessed with the Nanopolish 0.13.2 polya function^74^. For downstream analyses, only reads tagged by Nanopolish as PASS or SUFFCLIP were considered.

Initial processing of cDNA sequencing reads (basecalling, mapping, demultiplexing and poly(A) tail length estimation) were performed with Dorado (ONT) using default parameters. The lengths of poly(A) tails were then extracted from the source files and aggregated in tabular form. For the downstream analysis, only reads with mapping quality >0 were taken into account.

The poly(A) tail predictions from DRS and cDNA data were further processed in R environment. The NanoTail package^75^ was used to perform exploratory data analysis and statistical inference. Data were tested for normality using the Shapiro-Wilk test. The p-values were calculated using the Wilcoxon signed-rank test for transcripts represented by 10 or more reads and adjusted for multiple comparisons with the Benjamini-Hochberg method. Transcripts were considered significantly altered in poly(A) tail length if the adjusted p-value was less than 0.05 and the difference in tail length was greater than 5 nt between the conditions. The results were visualized in R using ggplot2 package.

### LC-MS/MS raw data analysis

The resulting RAW files were analyzed using MaxQuant^76,77^ version 2.0.3.0. Protein identification was performed against the the UniProt mouse (Mus musculus 10090) reference database (downloaded in July 2022), with trypsin-specific digestion and a maximum allowance of 2 missed cleavages. Variable modifications considered included methionine oxidation and N-terminal acetylation. A 1% false discovery rate (FDR) was applied to both peptide and protein identifications to ensure data quality.

### MS/MS processed data analysis

MaxLFQ Unique Intensity and Protein Length parameters returned by MAXQuant were used to calculate the enrichment (relative specificity) and abundance (relative intensity) of identified proteins in each biological replicate which was considered a pair of TENT5C-EGFP and WT erythroblasts subjected to TurboID modified protocol. First, all MaxLFQ Unique Intensity values which were equal to 0 were transformed to 1 to avoid dividing by 0. Relative specificity for each protein across all replicates was calculated by dividing transformed MaxLFQ Unique Intensity of TENT5C-EGFP by transformed MaxLFQ Unique Intensity of WT. Relative intensity was calculated by dividing transformed MaxLFQ Unique Intensity of TENT5C-EGFP by Protein Length. Next, filtering was applied to retain the values of relative specificity > 0 and overlap for 3 replicates was calculated. Proteins found in the overlap for 3 replicates were considered as enriched and this group was further analysed. GO enrichment analysis was performed with gprofiler2 package^78^ using default parameters and selected ontologies were visualized.

### TENT5C-EGFP depletion efficiency calculation

TENT5C-EGFP relative expression (Fig. S5C) was calculated as fold-change relative to control shRNA derived from gMFI of GFP channel in live singlets expressing mCherry diminished by autofluorescence of live WT singlets.

### TENT5C-EGFP half-lives calculation

Half-lives (Fig. S6D) were calculated based on an exponential decay model using cytometry data from CHX-chase experiments. TENT5C-EGFP gMFI diminished by autofluorescence of live WT singlets was used as protein level indicator. Measurements were taken at multiple time points following inhibition as described above. Relative protein levels were calculated for each condition with 0h timepoint as baseline. The natural logarithm of the relative protein levels was plotted against time, and a linear regression was performed. The decay constant (k) was obtained from the slope of the regression line. The half-life (t_1/2_) was then calculated using the formula: t_1/2_ = (ln(2)/-k).

### Degron prediction and TENT5C sequence conservation analysis

We used canonical uniprot sequence of murine TENT5C (Q5SSF7) to run following analyses. We used Degronopedia^79^ web server (https://degronopedia.com) for degron motif prediction. Evolutionary sequence conservation was determined using ConSurf^80^ web server (https://consurf.tau.ac.il). Both analyses were run with default parameters. The results of ConSurf analysis were visualized with ChimeraX v1.8^81^ as color coded evolutionary conservation score plotted on TENT5C alpha-fold 2 structure model. Predicted degron motifs with exposed residues were highlighted on structure model.

## RESOURCE AVAILABILITY

### Lead contact

Further information and requests for resources and reagents should be directed to and will be fulfilled by the lead contact, Andrzej Dziembowski.

### Materials availability

This study did not generate new unique reagents.

### Data and code availability

The raw DRS sequencing data (fast5 files) were deposited at the European Nucleotide Archive (ENA) (https://www.ebi.ac.uk/ena/browser/home) under the project numbers PRJEB75356 and PRJEB80123 with sample numbers listed in Key resource table. The cDNA sequencing data (fastq files) along with corresponding poly(A) length predictions and differential adenylation data are deposited at the Gene Expression Omnibus (GEO) (https://www.ncbi.nlm.nih.gov/geo/) under the project number PRJNA1177422 (https://www.ncbi.nlm.nih.gov/geo/query/acc.cgi?acc=GSE280237) with sample numbers listed in Key resource table. The raw MS/MS data was deposited to ZENODO (https://zenodo.org/records/14163383) alongside with any other supplementary data.

## ACKNOWLEDGEMENTS

We thank all members of the Laboratory of RNA Biology (LRB) and Genome Engineering Facility (GEU) at IIMCB for their expertise and help in selected experiments. We also thank Prof. H. Lodish for sharing retroviral vectors for shRNA expression. All mice lines were designed, generated and genotyped by the Genome Engineering Facility (GEU), part of IIMCB IN-MOL-CELL Infrastructure (RRID: SCR_021630) funded by the European Union – NextGenerationEU under National Recovery and Resilience Plan, Horizon Europe (Project 101059801 –RACE) and RACE-PRIME project carried out within the IRAP programme of the Foundation for Polish Science co-financed by the European Union under the European Funds for Smart Economy 2021-2027 (FENG) IIMCB.

This work was mainly supported by the National Science Center (SONATA 15 2019/35/D/NZ3/04253 to M.K-K.) and co-supported by (PRELUDIUM 19 2020/37/N/NZ2/02893 to A.B.) and HORIZON Europa (ERC AdG 101097317 to A.D.).

## AUTHOR CONTRIBUTIONS

A.D. and M.K-K. conceived the project; A.D., M.K-K. and M.M. designed the study; M.K-K. conducted phenotyping experiments and analyzed related data; M.M. conducted the majority of experiments, analyzed related data, and conceptualized the research; A.B. validated TENT5C-LARP4/5 interaction in HEK293T model; N.G. analyzed all sequencing data, prepared related figures and ensured graphical integrity of all figures; D.C processed TurboID samples after pulldown and analyzed the raw data; K.M-S. and M.N. did iron measurement experiments. N.G., A.B., K.M-S. provided feedback on the manuscript. M.M., A.D., and M.K-K. wrote the manuscript, with contributions from other authors. All authors read and approved the manuscript.

## FIGURES

**Supplementary figure S1.**
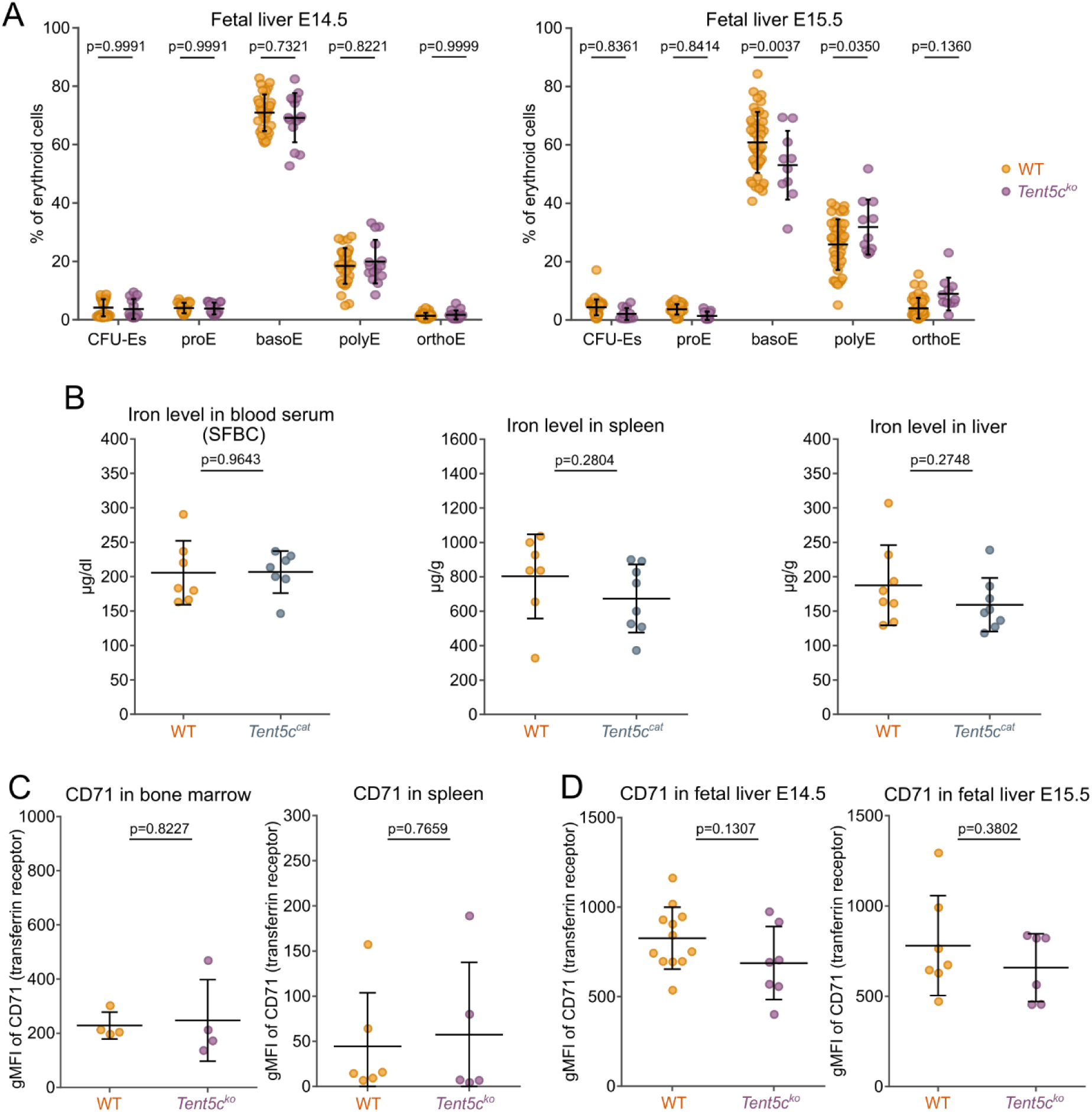
TENT5C-dependent anemic phenotype is not related to iron pathways. (A) Erythroblasts differentiation in fetal liver in E14.5 (left panel) and E15.5 (right panel) (WT *n*=20, *Tent5c^ko^ n*=11-14). Data presents as mean ± SD, and significance assessed by two-way ANOVA. (B) Iron level in blood serum (left panel), spleen (middle panel) and liver (right panel) (WT *n*=7, *Tent5c^cat^ n*=7). Data presented as mean ± SD, and significance was assessed by *t*-Student test. (C) Quantification (gMFI) of CD71 expression erythroid cells isolated from the bone marrow (left panel) and the spleen (right panel) (WT *n*=4-6, *Tent5c^ko^ n*=4-6). Data presented as mean ± SD, and significance was assessed by *t*-Student test. (D) Quantification (gMFI) of CD71 expression in erythroid cells isolated from the fetal livers at E14.5 (left panel) and E15.5 (right panel) embryonic development days (WT *n*=7-12, *Tent5c^ko^ n*=6-7). Data presented as mean ± SD, and significance was assessed by *t*-Student test.

**Supplementary figure S2.**
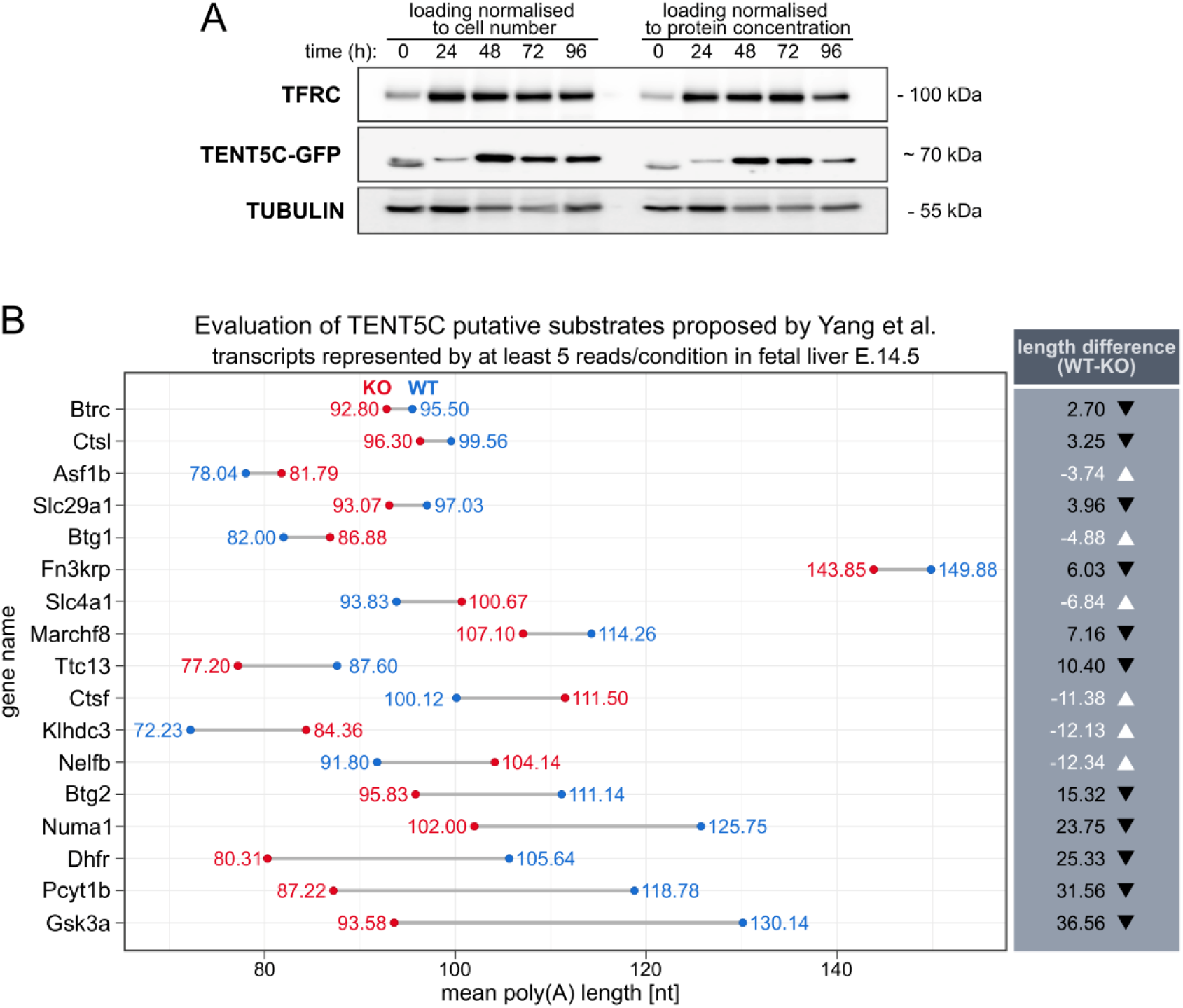
TENT5C expression *ex vivo* and evaluation of putative TENT5C substrates suggested by Yang et al. (A) Western Blot representing TENT5C-EGFP protein expression in time course of *Tent5c-EGFP* murine erythroblasts cultured *ex vivo*. The left side is loaded according to cell number, the right side is loaded according to lysate protein concentration. 0h timepoint corresponds to isolated progenitors used to start the culture. (B) The graph is based on DRS data from E14.5 fetal liver. Only genes represented by at least five reads per condition are included. The differences in mean poly(A) tail lengths are shown. Genes are sorted from top to bottom according to increasing absolute difference values, with WT marked in blue and KO marked in red. The right panel displays the WT-KO difference values. Black text with a downward-pointing arrow denotes poly(A) tail shortening in KO, while white text with an upward-pointing arrow indicates poly(A) tail lengthening in KO.

**Supplementary figure S3.**
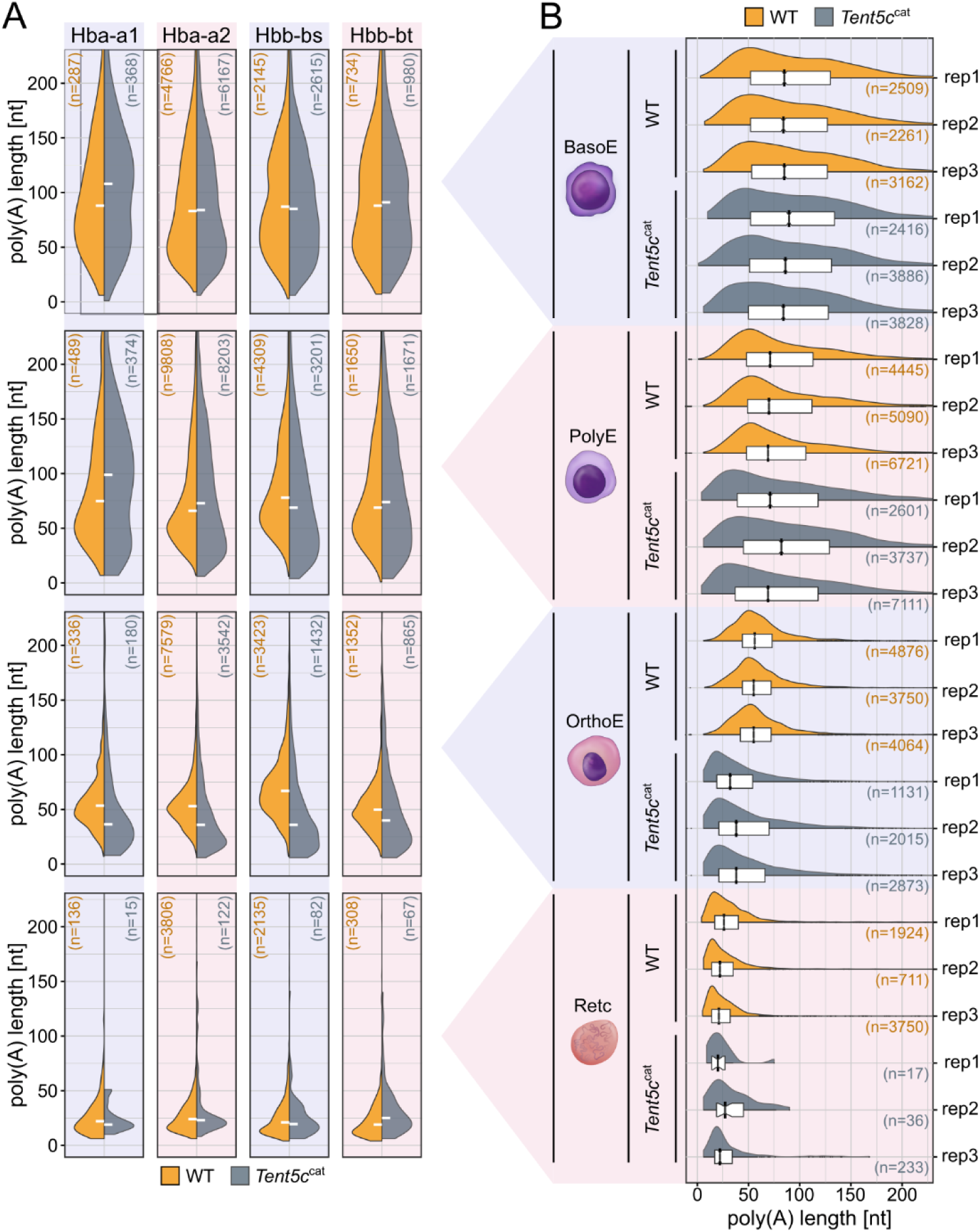
Globin expression pattern during erythropoiesis. (A) Poly(A) tail lengths distribution in each globin transcript during erythropoiesis in WT and *Tent5c^cat^*. Data grouped by condition. White horizontal lines indicate medians. Numbers of reads in each group (n) are shown above the distribution plots. (B) Poly(A) tail lengths distribution for combined globin reads during erythropoiesis in WT and *Tent5c^cat^*. Data grouped by replicate. Black vertical lines represent the median, while the lower and upper hinges correspond to the first and third quartiles (25th and 75th percentiles). The whiskers extend to the smallest and largest values within 1.5 times the interquartile range from the hinges. Numbers of reads in each group (n) are shown above the distribution plots.

**Supplementary figure S4.**
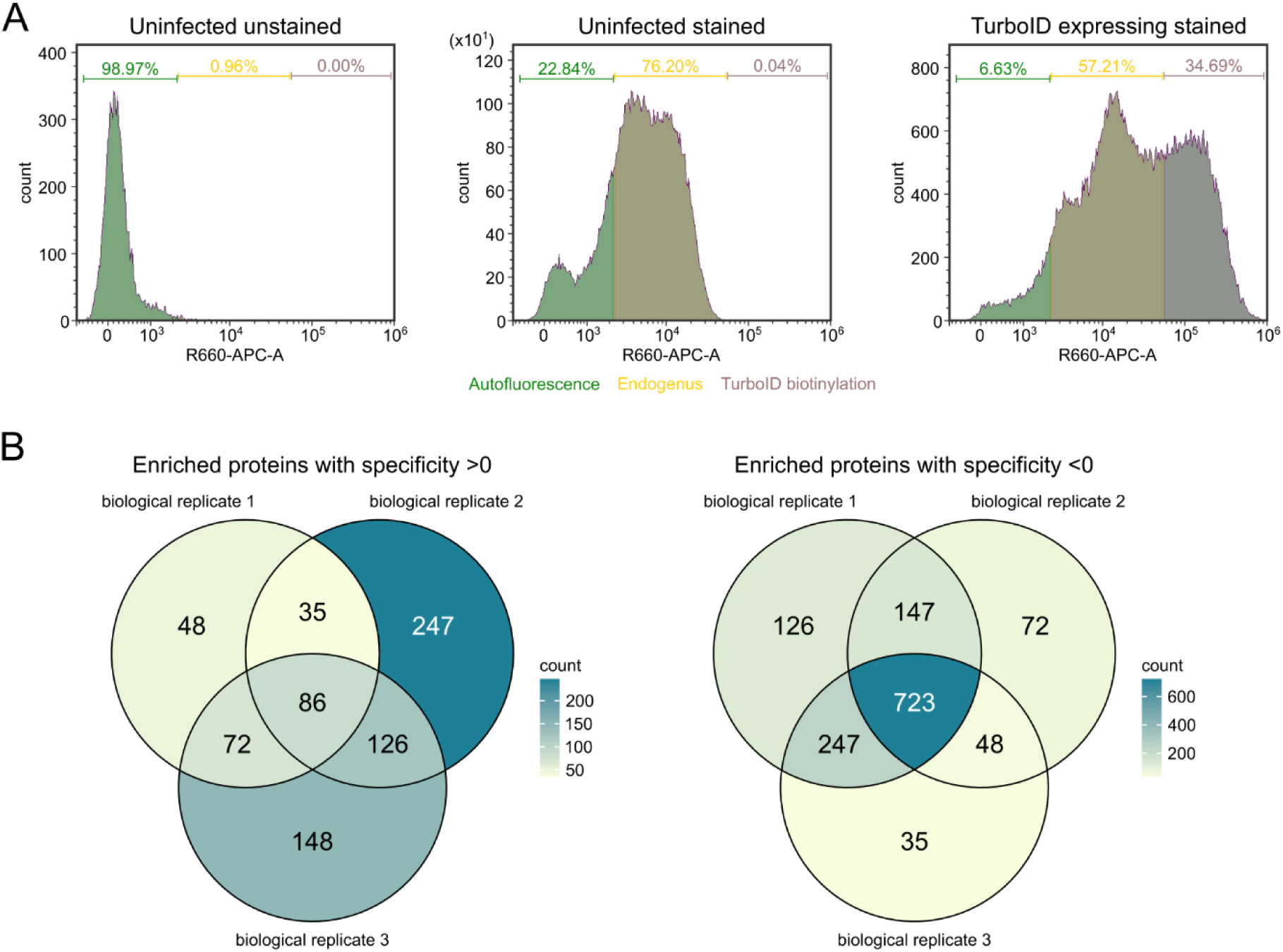
TurboID supplementary information. (A) Histograms representing intracellular staining of erythroblasts from TurboID experiment with Streptavidin-AlexaFluor 647. Samples arranged (left to right): uninfected unstained, uninfected stained, infected stained. (B) Venn plots representing enriched (left) and background (right) proteins across 3 biological replicates of TurboID experiment.

**Supplementary figure S5.**
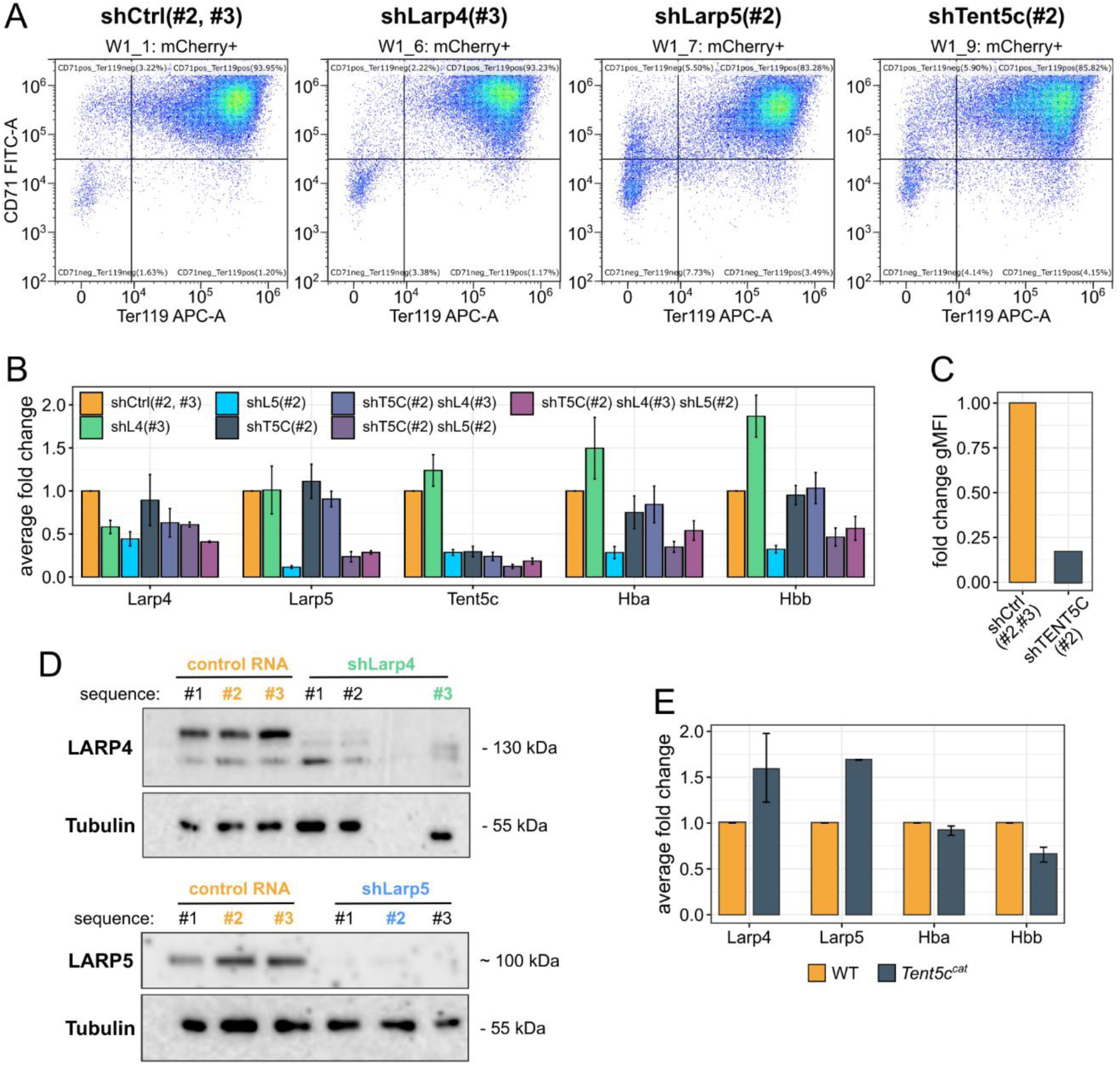
Supplementary data on shRNA silencing. (A) Flow cytometry dot-plots representing the major trajectory of erythroblast differentiation in cells expressing shRNA (control, shLarp4, Larp5, Tent5c) with respect to CD71 (y-axis) and TER119 (x-axis). (B) RT-qPCR results for selected genes in samples with different depletions (n=3). Average fold changes relative to shCtrl (#2, #3) with *Gapdh* expression used for normalization. Error bars correspond to the standard error of the mean (SEM). (C) TENT5C protein expression in cells expressing shRNA against Tent5c vs shCtrl(#2, #3). Expression based on gMFI, calculated as a fold-change relative to control. (D) Western Blot depicting LARP4 (top) or LARP5 (bottom) depletion efficiency compared to control shRNAs. Cells acquired by FACS; equal cell number loaded onto wells (25000 cells). Selected shRNA used in experiments were highlighted in bold and color. (E) RT-qPCR results for Larp4/5 and globin genes in WT and *Tent5c^cat^* erythroblasts (mixed population) from the bone marrow (n=2). Average fold changes relative to WT with *Gapdh* expression used as a normalization factor. Error bars correspond to the standard error of the mean (SEM).

**Supplementary figure S6.**
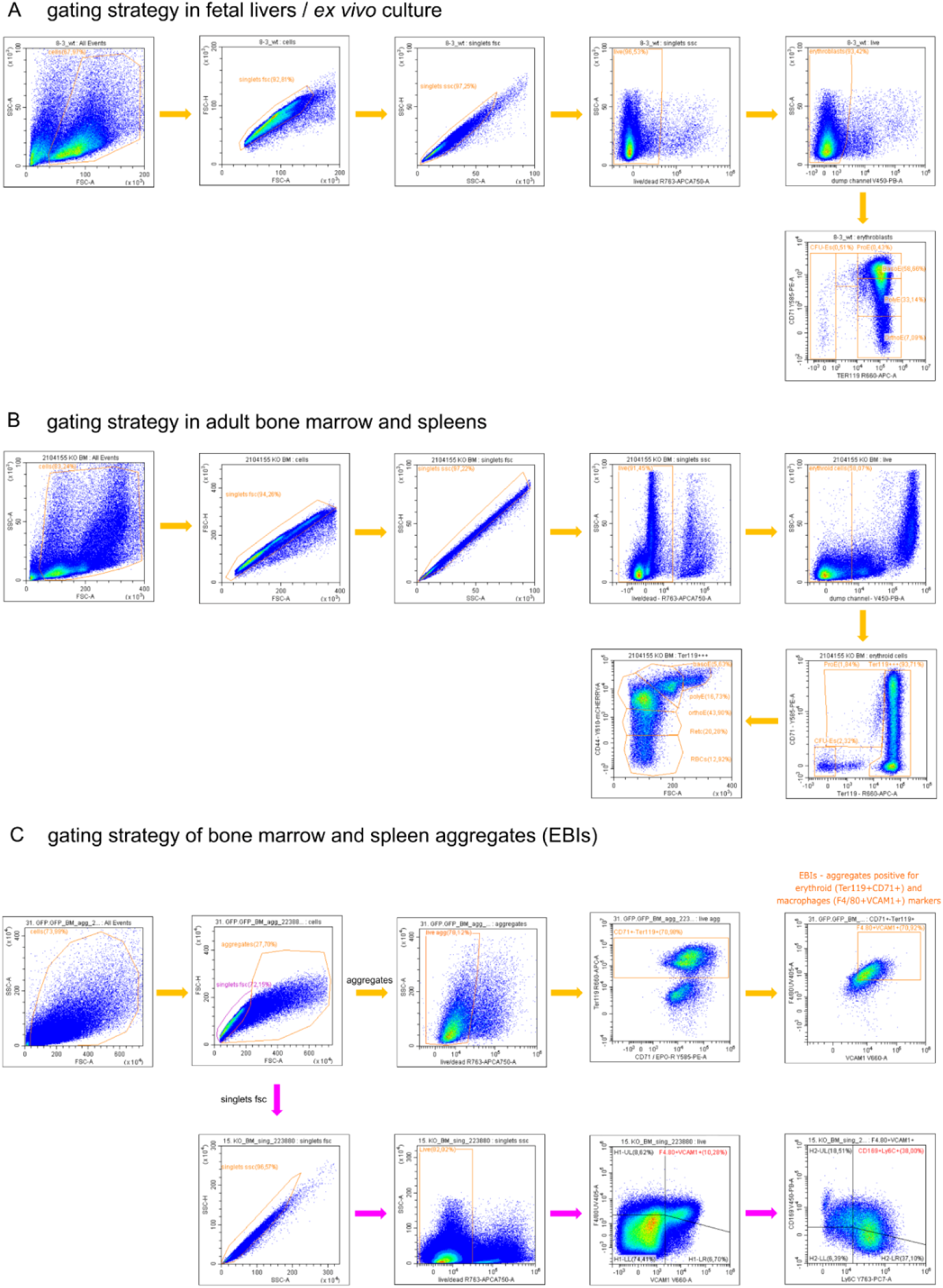
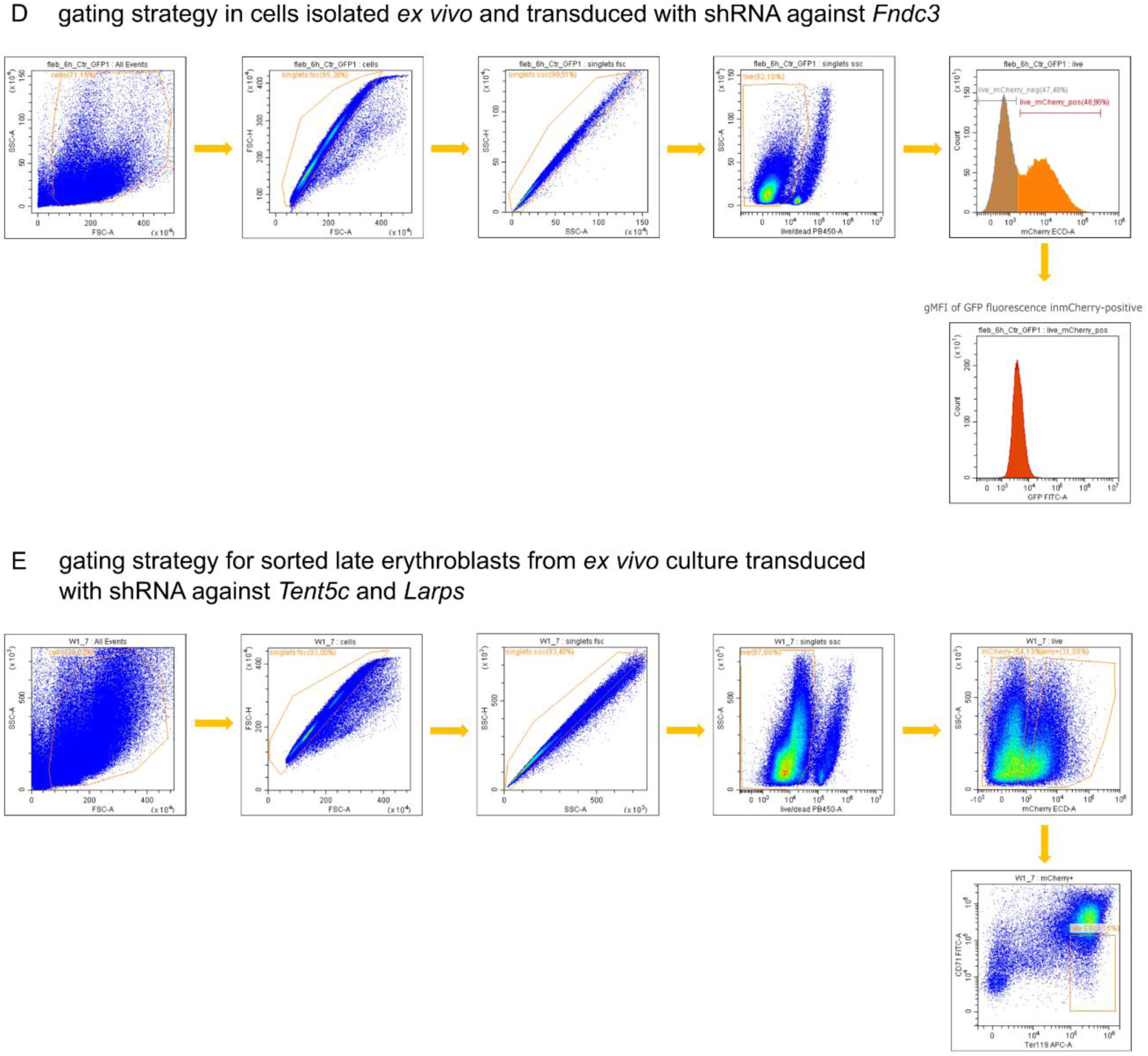
Gating strategy used in experiments. (A) Gating strategy used for cells isolated from fetal livers and *ex vivo* cultures. (B) Gating strategy used for cells isolated from adult bone marrow and spleen. (C) Gating strategy used for EBIs calculation and characterization of EBIs macrophages. (D) Gating strategy used for cells isolated ex vivo and transduced with shRNA against *Cnot4*. (E) Gating strategy for sorted late erythroblasts from ex vivo culture transduced with shRNA against *Tent5c* and *Larps*.

## REFERENCES

1. Ravenhill, B.J., Kanjee, U., Ahouidi, A., Nobre, L., Williamson, J., Goldberg, J.M., Antrobus, R., Dieye, T., Duraisingh, M.T., and Weekes, M.P. (2019). Quantitative comparative analysis of human erythrocyte surface proteins between individuals from two genetically distinct populations. Commun. Biol. 2, 350. 10.1038/s42003-019-0596-y.

2. Roux-Dalvai, F., Gonzalez De Peredo, A., Simoé, C., Guerrier, L., Bouyssieé, D., Zanella, A., Citterio, A., Burlet-Schiltz, O., Boschetti, E., Righetti, P.G., et al. (2008). Extensive Analysis of the Cytoplasmic Proteome of Human Erythrocytes Using the Peptide Ligand Library Technology and Advanced Mass Spectrometry. Mol. Cell. Proteomics 7, 2254–2269. 10.1074/mcp.M800037-MCP200.

3. Gautier, E.-F., Ducamp, S., Leduc, M., Salnot, V., Guillonneau, F., Dussiot, M., Hale, J., Giarratana, M.-C., Raimbault, A., Douay, L., et al. (2016). Comprehensive Proteomic Analysis of Human Erythropoiesis. Cell Rep. 16, 1470–1484. 10.1016/j.celrep.2016.06.085.

4. Gautier, E.-F., Leduc, M., Ladli, M., Schulz, V.P., Lefèvre, C., Boussaid, I., Fontenay, M., Lacombe, C., Verdier, F., Guillonneau, F., et al. (2020). Comprehensive proteomic analysis of murine terminal erythroid differentiation. Blood Adv. 4, 1464–1477. 10.1182/bloodadvances.2020001652.

5. Chen, K., Liu, J., Heck, S., Chasis, J.A., An, X., and Mohandas, N. (2009). Resolving the distinct stages in erythroid differentiation based on dynamic changes in membrane protein expression during erythropoiesis. Proc. Natl. Acad. Sci. 106, 17413–17418. 10.1073/pnas.0909296106.

6. Moras, M., Lefevre, S.D., and Ostuni, M.A. (2017). From Erythroblasts to Mature Red Blood Cells: Organelle Clearance in Mammals. Front. Physiol. 8, 1076. 10.3389/fphys.2017.01076.

7. Dzierzak, E., and Philipsen, S. (2013). Erythropoiesis: Development and Differentiation. Cold Spring Harb. Perspect. Med. 3, a011601–a011601. 10.1101/cshperspect.a011601.

8. McGrath, K.E., Catherman, S.C., and Palis, J. (2017). Delineating stages of erythropoiesis using imaging flow cytometry. Methods 112, 68–74. 10.1016/j.ymeth.2016.08.012.

9. Liao, C., Prabhu, K.S., and Paulson, R.F. (2018). Monocyte-derived macrophages expand the murine stress erythropoietic niche during the recovery from anemia. Blood 132, 2580–2593. 10.1182/blood-2018-06-856831.

10. Seu, K.G., Papoin, J., Fessler, R., Hom, J., Huang, G., Mohandas, N., Blanc, L., and Kalfa, T.A. (2017). Unraveling Macrophage Heterogeneity in Erythroblastic Islands. Front. Immunol. 8, 1140. 10.3389/fimmu.2017.01140.

11. Li, W., Guo, R., Song, Y., and Jiang, Z. (2021). Erythroblastic Island Macrophages Shape Normal Erythropoiesis and Drive Associated Disorders in Erythroid Hematopoietic Diseases. Front. Cell Dev. Biol. 8, 613885. 10.3389/fcell.2020.613885.

12. Ji, P. (2015). New Insights into the Mechanisms of Mammalian Erythroid Chromatin Condensation and Enucleation. In International Review of Cell and Molecular Biology (Elsevier), pp. 159–182. 10.1016/bs.ircmb.2015.01.006.

13. Koulnis, M., Pop, R., Porpiglia, E., Shearstone, J.R., Hidalgo, D., and Socolovsky, M. (2011). Identification and Analysis of Mouse Erythroid Progenitors using the CD71/TER119 Flow-cytometric Assay. J. Vis. Exp., 2809. 10.3791/2809.

14. Palis, J., and Segel, G.B. (1998). Developmental biology of erythropoiesis. Blood Rev. 12, 106– 114. 10.1016/S0268-960X(98)90022-4.

15. Palis, J. (2014). Primitive and definitive erythropoiesis in mammals. Front. Physiol. 5. 10.3389/fphys.2014.00003.

16. Zhao, B., Mei, Y., Yang, J., and Ji, P. (2014). Mouse fetal liver culture system to dissect target gene functions at the early and late stages of terminal erythropoiesis. J. Vis. Exp. JoVE, e51894. 10.3791/51894.

17. Bennett, L.F., Liao, C., Quickel, M.D., Yeoh, B.S., Vijay-Kumar, M., Hankey-Giblin, P., Prabhu, K.S., and Paulson, R.F. (2019). Inflammation induces stress erythropoiesis through heme-dependent activation of SPI-C. Sci. Signal. 12, eaap7336. 10.1126/scisignal.aap7336.

18. Hattangadi, S.M., Wong, P., Zhang, L., Flygare, J., and Lodish, H.F. (2011). From stem cell to red cell: regulation of erythropoiesis at multiple levels by multiple proteins, RNAs, and chromatin modifications. Blood 118, 6258–6268. 10.1182/blood-2011-07-356006.

19. Udroiu, I. (2016). Development of erythropoiesis in the mouse. Russ. J. Dev. Biol. 47, 254–259. 10.1134/S1062360416050052.

20. Paulson, R.F., Shi, L., and Wu, D.-C. (2011). Stress erythropoiesis: new signals and new stress progenitor cells: Curr. Opin. Hematol. 18, 139–145. 10.1097/MOH.0b013e32834521c8.

21. Bennett, L.F., Liao, C., and Paulson, R.F. (2018). Stress Erythropoiesis Model Systems. In Erythropoiesis, J. A. Lloyd, ed. (Springer New York), pp. 91–102. 10.1007/978-1-4939-7428-3_5.

22. Socolovsky, M. (2007). Molecular insights into stress erythropoiesis: Curr. Opin. Hematol. 14, 215–224. 10.1097/MOH.0b013e3280de2bf1.

23. McCranor, B.J., Kim, M.J., Cruz, N.M., Xue, Q.-L., Berger, A.E., Walston, J.D., Civin, C.I., and Roy, C.N. (2014). Interleukin-6 directly impairs the erythroid development of human TF-1 erythroleukemic cells. Blood Cells. Mol. Dis. 52, 126–133. 10.1016/j.bcmd.2013.09.004.

24. Manchinu, M.F., Brancia, C., Caria, C.A., Musu, E., Porcu, S., Simbula, M., Asunis, I., Perseu, L., and Ristaldi, M.S. (2018). Deficiency in interferon type 1 receptor improves definitive erythropoiesis in Klf1 null mice. Cell Death Differ. 25, 589–599. 10.1038/s41418-017-0003-5.

25. Zermati, Y., Fichelson, S., Valensi, F., Freyssinier, J.M., Rouyer-Fessard, P., Cramer, E., Guichard, J., Varet, B., and Hermine, O. (2000). Transforming growth factor inhibits erythropoiesis by blocking proliferation and accelerating differentiation of erythroid progenitors. Exp. Hematol. 28, 885–894. 10.1016/S0301-472X(00)00488-4.

26. Han, G., Li, F., Singh, T.P., Wolf, P., and Wang, X.-J. (2012). The Pro-inflammatory Role of TGFβ1: A Paradox? Int. J. Biol. Sci. 8, 228–235. 10.7150/ijbs.8.228.

27. Murphy, Z.C., Murphy, K., Myers, J., Getman, M., Couch, T., Schulz, V.P., Lezon-Geyda, K., Palumbo, C., Yan, H., Mohandas, N., et al. (2021). Regulation of RNA polymerase II activity is essential for terminal erythroid maturation. Blood 138, 1740–1756. 10.1182/blood.2020009903.

28. An, X., Schulz, V.P., Li, J., Wu, K., Liu, J., Xue, F., Hu, J., Mohandas, N., and Gallagher, P.G. (2014). Global transcriptome analyses of human and murine terminal erythroid differentiation. Blood 123, 3466–3477. 10.1182/blood-2014-01-548305.

29. Nguyen, A.T., Prado, M.A., Schmidt, P.J., Sendamarai, A.K., Wilson-Grady, J.T., Min, M., Campagna, D.R., Tian, G., Shi, Y., Dederer, V., et al. (2017). UBE2O remodels the proteome during terminal erythroid differentiation. Science 357, eaan0218. 10.1126/science.aan0218.

30. Grosso, R., Fader, C.M., and Colombo, M.I. (2017). Autophagy: A necessary event during erythropoiesis. Blood Rev. 31, 300–305. 10.1016/j.blre.2017.04.001.

31. Chen, J.J., and London, I.M. (1995). Regulation of protein synthesis by heme-regulated eIF-2 alpha kinase. Trends Biochem. Sci. 20, 105–108. 10.1016/s0968-0004(00)88975-6.

32. Hollenberg, M.D., Kaback, M.M., and Kazazian, H.H. (1971). Adult hemoglobin synthesis by reticulocytes from the human fetus at midtrimester. Science 174, 698–702. 10.1126/science.174.4010.698.

33. Crosby, J.S., Chefalo, P.J., Yeh, I., Ying, S., London, I.M., Leboulch, P., and Chen, J.-J. (2000). Regulation of hemoglobin synthesis and proliferation of differentiating erythroid cells by heme-regulated eIF-2α kinase. Blood 96, 3241–3248. 10.1182/blood.V96.9.3241.

34. 34. Jorge, S.E., Ribeiro, D.M., Santos, M.N.N., and de Fátima Sonati, M. (2016). Hemoglobin: Structure, Synthesis and Oxygen Transport. In Sickle Cell Anemia: From Basic Science to Clinical Practice, F. F. Costa and N. Conran, eds. (Springer International Publishing), pp. 1–22. 10.1007/978-3-319-06713-1_1.

35. McIver, S.C., Kang, Y.-A., DeVilbiss, A.W., O’Driscoll, C.A., Ouellette, J.N., Pope, N.J., Camprecios, G., Chang, C.-J., Yang, D., Bouhassira, E.E., et al. (2014). The exosome complex establishes a barricade to erythroid maturation. Blood 124, 2285–2297. 10.1182/blood-2014-04-571083.

36. 36. Han, A., Yermalovich, A.V., Lundin, V., Pearson, D.S., Hachimi, M., Sousa, P., Hilton, B., Morse, M., Atwater, J., Zhang, Y., et al. (2020). An Essential Role for the RNA Editor-Exonuclease Axis in Terminal Erythroid Differentiation. Blood 136, 3. 10.1182/blood-2020-142757.

37. Waggoner, S.A., and Liebhaber, S.A. (2003). Regulation of α-Globin mRNA Stability. Exp. Biol. Med. 228, 387–395. 10.1177/153537020322800409.

38. Wang, Z., and Kiledjian, M. (2000). The Poly(A)-Binding Protein and an mRNA Stability Protein Jointly Regulate an Endoribonuclease Activity. Mol. Cell. Biol. 20, 6334. 10.1128/mcb.20.17.6334-6341.2000.

39. Mroczek, S., Chlebowska, J., Kuliński, T.M., Gewartowska, O., Gruchota, J., Cysewski, D., Liudkovska, V., Borsuk, E., Nowis, D., and Dziembowski, A. (2017). The non-canonical poly(A) polymerase FAM46C acts as an onco-suppressor in multiple myeloma. Nat. Commun. 8, 619. 10.1038/s41467-017-00578-5.

40. Cao, W., Fan, W., Wang, F., Zhang, Y., Wu, G., Shi, X., Shi, J.X., Gao, F., Yan, M., Guo, R., et al. (2022). GM-CSF impairs erythropoiesis by disrupting erythroblastic island formation via macrophages. J. Transl. Med. 20, 11. 10.1186/s12967-021-03214-5.

41. Brouze, A., Krawczyk, P.S., Dziembowski, A., and Mroczek, S. (2023). Measuring the tail: Methods for poly(A) tail profiling. WIREs RNA 14, e1737. 10.1002/wrna.1737.

42. Bilska, A., Kusio-Kobiałka, M., Krawczyk, P.S., Gewartowska, O., Tarkowski, B., Kobyłecki, K., Nowis, D., Golab, J., Gruchota, J., Borsuk, E., et al. (2020). Immunoglobulin expression and the humoral immune response is regulated by the non-canonical poly(A) polymerase TENT5C. Nat. Commun. 11, 2032. 10.1038/s41467-020-15835-3.

43. Liudkovska, V., Krawczyk, P.S., Brouze, A., Gumińska, N., Wegierski, T., Cysewski, D., Mackiewicz, Z., Ewbank, J.J., Drabikowski, K., Mroczek, S., et al. (2022). TENT5 cytoplasmic noncanonical poly(A) polymerases regulate the innate immune response in animals. Sci. Adv. 8, eadd9468. 10.1126/sciadv.add9468.

44. Gewartowska, O., Aranaz-Novaliches, G., Krawczyk, P.S., Mroczek, S., Kusio-Kobiałka, M., Tarkowski, B., Spoutil, F., Benada, O., Kofroňová, O., Szwedziak, P., et al. (2021). Cytoplasmic polyadenylation by TENT5A is required for proper bone formation. Cell Rep. 35, 109015. 10.1016/j.celrep.2021.109015.

45. Brouze, M., Czarnocka-Cieciura, A., Gewartowska, O., Kusio-Kobiałka, M., Jachacy, K., Szpila, M., Tarkowski, B., Gruchota, J., Krawczyk, P., Mroczek, S., et al. (2024). TENT5-mediated polyadenylation of mRNAs encoding secreted proteins is essential for gametogenesis in mice. Nat. Commun. 15, 5331. 10.1038/s41467-024-49479-4.

46. Yang, K., Zhu, T., Yin, J., Zhang, Q., Li, J., Fan, H., Han, G., Xu, W., Liu, N., and Lv, X. (2024). The non-canonical poly(A) polymerase FAM46C promotes erythropoiesis. J. Genet. Genomics Yi Chuan Xue Bao, S1673–8527(24)00032-8. 10.1016/j.jgg.2024.02.003.

47. Muhlrad, D., Decker, C.J., and Parker, R. (1994). Deadenylation of the unstable mRNA encoded by the yeast MFA2 gene leads to decapping followed by 5’-->3’ digestion of the transcript. Genes Dev. 8, 855–866. 10.1101/gad.8.7.855.

48. Decker, C.J., and Parker, R. (1993). A turnover pathway for both stable and unstable mRNAs in yeast: evidence for a requirement for deadenylation. Genes Dev. 7, 1632–1643. 10.1101/gad.7.8.1632.

49. Fucci, C., Resnati, M., Riva, E., Perini, T., Ruggieri, E., Orfanelli, U., Paradiso, F., Cremasco, F., Raimondi, A., Pasqualetto, E., et al. (2020). The Interaction of the Tumor Suppressor FAM46C with p62 and FNDC3 Proteins Integrates Protein and Secretory Homeostasis. Cell Rep. 32, 108162. 10.1016/j.celrep.2020.108162.

50. Branon, T.C., Bosch, J.A., Sanchez, A.D., Udeshi, N.D., Svinkina, T., Carr, S.A., Feldman, J.L., Perrimon, N., and Ting, A.Y. (2018). Efficient proximity labeling in living cells and organisms with TurboID. Nat. Biotechnol. 36, 880–887. 10.1038/nbt.4201.

51. 51. Tian, M. (2010). The molecular cloning and characterization of Fam46c RNA stability factor.

52. Russell, J.E., and Liebhaber, S.A. (1996). The stability of human beta-globin mRNA is dependent on structural determinants positioned within its 3’ untranslated region. Blood 87, 5314–5323.

53. Russell, J.E., Morales, J., and Liebhaber, S.A. (1997). The role of mRNA stability in the control of globin gene expression. Prog. Nucleic Acid Res. Mol. Biol. 57, 249–287. 10.1016/s0079-6603(08)60283-4.

54. Thein, S.L. (2013). The Molecular Basis of β-Thalassemia. Cold Spring Harb. Perspect. Med. 3, a011700. 10.1101/cshperspect.a011700.

55. 55. Keohane, E.M. (2020). 25 –Thalassemias. In Rodak’s Hematology (Sixth Edition), E. M. Keohane, C. N. Otto, and J. M. Walenga, eds. (Elsevier), pp. 424–444. 10.1016/B978-0-323-53045-3.00034-9.

56. Collart, M.A. (2016). The Ccr4-Not complex is a key regulator of eukaryotic gene expression. Wiley Interdiscip. Rev. RNA 7, 438. 10.1002/wrna.1332.

57. Buschauer, R., Matsuo, Y., Sugiyama, T., Chen, Y.-H., Alhusaini, N., Sweet, T., Ikeuchi, K., Cheng, J., Matsuki, Y., Nobuta, R., et al. (2020). The Ccr4-Not complex monitors the translating ribosome for codon optimality. Science 368, eaay6912. 10.1126/science.aay6912.

58. Maraia, R.J., Mattijssen, S., Cruz-Gallardo, I., and Conte, M.R. (2017). The LARPs, La and related RNA-binding proteins: Structures, functions and evolving perspectives. Wiley Interdiscip. Rev. RNA 8, 10.1002/wrna.1430. 10.1002/wrna.1430.

59. Deragon, J.-M. (2021). Distribution, organization an evolutionary history of La and LARPs in eukaryotes. RNA Biol. 18, 159–167. 10.1080/15476286.2020.1739930.

60. Mattijssen, S., Kozlov, G., Fonseca, B.D., Gehring, K., and Maraia, R.J. (2021). LARP1 and LARP4: up close with PABP for mRNA 3’ poly(A) protection and stabilization. RNA Biol. 18, 259–274. 10.1080/15476286.2020.1868753.

61. 61. Cruz-Gallardo, I., Martino, L., Kelly, G., Atkinson, R.A., Trotta, R., De Tito, S., Coleman, P., Ahdash, Z., Gu, Y., Bui, T.T.T., et al. (2019). LARP4A recognizes polyA RNA via a novel binding mechanism mediated by disordered regions and involving the PAM2w motif, revealing interplay between PABP, LARP4A and mRNA. Nucleic Acids Res. 47, 4272–4291. 10.1093/nar/gkz144.

62. Schäffler, K., Schulz, K., Hirmer, A., Wiesner, J., Grimm, M., Sickmann, A., and Fischer, U. (2010). A stimulatory role for the La-related protein 4B in translation. RNA N. Y. N 16, 1488–1499. 10.1261/rna.2146910.

63. Küspert, M., Murakawa, Y., Schäffler, K., Vanselow, J.T., Wolf, E., Juranek, S., Schlosser, A., Landthaler, M., and Fischer, U. (2015). LARP4B is an AU-rich sequence associated factor that promotes mRNA accumulation and translation. RNA 21, 1294. 10.1261/rna.051441.115.

64. Tay, J., Bisht, K., McGirr, C., Millard, S.M., Pettit, A.R., Winkler, I.G., and Levesque, J.-P. (2020). Imaging flow cytometry reveals that granulocyte colony-stimulating factor treatment causes loss of erythroblastic islands in the mouse bone marrow. Exp. Hematol. 82, 33–42. 10.1016/j.exphem.2020.02.003.

65. Fujiyama, S., Nakahashi-Oda, C., Abe, F., Wang, Y., Sato, K., and Shibuya, A. (2019). Identification and isolation of splenic tissue-resident macrophage sub-populations by flow cytometry. Int. Immunol. 31, 51–56. 10.1093/intimm/dxy064.

66. Duarte, T.L., Jo&#227, and Neves, o V. (2022). Measurement of Tissue Non-Heme Iron Content using a Bathophenanthroline-Based Colorimetric Assay. JoVE J. Vis. Exp., e63469. 10.3791/63469.

67. Bajak, E.Z., and Hagedorn, C.H. (2008). Efficient 5’ cap-dependent RNA purification : use in identifying and studying subsets of RNA. Methods Mol. Biol. Clifton NJ 419, 147–160. 10.1007/978-1-59745-033-1_10.

68. Choi, Y.H., and Hagedorn, C.H. (2003). Purifying mRNAs with a high-affinity eIF4E mutant identifies the short 3’ poly(A) end phenotype. Proc. Natl. Acad. Sci. U. S. A. 100, 7033–7038. 10.1073/pnas.1232347100.

69. Ye, J., Coulouris, G., Zaretskaya, I., Cutcutache, I., Rozen, S., and Madden, T.L. (2012). Primer-BLAST: A tool to design target-specific primers for polymerase chain reaction. BMC Bioinformatics 13, 134. 10.1186/1471-2105-13-134.

70. Livak, K.J., and Schmittgen, T.D. (2001). Analysis of relative gene expression data using real-time quantitative PCR and the 2(-Delta Delta C(T)) Method. Methods San Diego Calif 25, 402–408. 10.1006/meth.2001.1262.

71. Hughes, C.S., Moggridge, S., Müller, T., Sorensen, P.H., Morin, G.B., and Krijgsveld, J. (2019). Single-pot, solid-phase-enhanced sample preparation for proteomics experiments. Nat. Protoc. 14, 68–85. 10.1038/s41596-018-0082-x.

72. Li, H. (2018). Minimap2: pairwise alignment for nucleotide sequences. Bioinformatics 34, 3094– 3100. 10.1093/bioinformatics/bty191.

73. Danecek, P., Bonfield, J.K., Liddle, J., Marshall, J., Ohan, V., Pollard, M.O., Whitwham, A., Keane, T., McCarthy, S.A., Davies, R.M., et al. (2021). Twelve years of SAMtools and BCFtools. GigaScience 10, giab008. 10.1093/gigascience/giab008.

74. Workman, R.E., Tang, A.D., Tang, P.S., Jain, M., Tyson, J.R., Razaghi, R., Zuzarte, P.C., Gilpatrick, T., Payne, A., Quick, J., et al. (2019). Nanopore native RNA sequencing of a human poly(A) transcriptome. Nat. Methods 16, 1297–1305. 10.1038/s41592-019-0617-2.

75. Krawczyk, P. (2024). NanoTail –R package for exploratory analysis of Nanopore Direct RNA based polyA lengths estimations.

76. Cox, J., and Mann, M. (2008). MaxQuant enables high peptide identification rates, individualized p.p.b.-range mass accuracies and proteome-wide protein quantification. Nat. Biotechnol. 26, 1367–1372. 10.1038/nbt.1511.

77. Cox, J., Michalski, A., and Mann, M. (2011). Software Lock Mass by Two-Dimensional Minimization of Peptide Mass Errors. J. Am. Soc. Mass Spectrom. 22, 1373–1380. 10.1007/s13361-011-0142-8.

78. Kolberg, L., Raudvere, U., Kuzmin, I., Vilo, J., and Peterson, H. (2020). gprofiler2 –-an R package for gene list functional enrichment analysis and namespace conversion toolset g:Profiler. F1000Research 9, ELIXIR. 10.12688/f1000research.24956.2.

79. Szulc, N.A., Stefaniak, F., Piechota, M., Soszyńska, A., Piórkowska, G., Cappannini, A., Bujnicki, J.M., Maniaci, C., and Pokrzywa, W. (2024). DEGRONOPEDIA: a web server for proteome-wide inspection of degrons. Nucleic Acids Res. 52, W221–W232. 10.1093/nar/gkae238.

80. Ashkenazy, H., Abadi, S., Martz, E., Chay, O., Mayrose, I., Pupko, T., and Ben-Tal, N. (2016). ConSurf 2016: an improved methodology to estimate and visualize evolutionary conservation in macromolecules. Nucleic Acids Res. 44, W344–350. 10.1093/nar/gkw408.

81. Meng, E.C., Goddard, T.D., Pettersen, E.F., Couch, G.S., Pearson, Z.J., Morris, J.H., and Ferrin, T.E. (2023). UCSF ChimeraX: Tools for structure building and analysis. Protein Sci. 32, e4792. 10.1002/pro.4792.

